# IGF-1 and insulin receptors in LepRb neurons jointly regulate body growth, bone mass, reproduction, and metabolism

**DOI:** 10.1101/2024.09.20.614140

**Authors:** Mengjie Wang, Piotr J. Czernik, Beata Lecka-Czernik, Yong Xu, Jennifer W. Hill

## Abstract

Leptin receptor (LepRb)-expressing neurons are known to link body growth and reproduction, but whether these functions are mediated via insulin-like growth factor 1 receptor (IGF1R) signaling is unknown. IGF-1 and insulin can bind to each other’s receptors, permitting IGF-1 signaling in the absence of IGF1R. Therefore, we created mice lacking IGF1R exclusively in LepRb neurons (IGF1R^LepRb^ mice) and simultaneously lacking IGF1R and insulin receptor (IR) in LepRb neurons (IGF1R/IR^LepRb^ mice) and then characterized their body growth, bone morphology, reproductive and metabolic functions. We found that IGF1R and IR in LepRb neurons were required for normal timing of pubertal onset, while IGF1R in LepRb neurons played a predominant role in regulating adult fertility and exerted protective effects against reproductive aging. Accompanying these reproductive deficits, IGF1R^LepRb^ mice and IGF1R/IR^LepRb^ mice had transient growth retardation. Notably, IGF1R in LepRb neurons was indispensable for normal trabecular and cortical bone mass accrual in both sexes. These findings suggest that IGF1R in LepRb neurons is involved in the interaction among body growth, bone development, and reproduction. Though only mild changes in body weight were detected, simultaneous deletion of IGF1R and IR in LepRb neurons caused dramatically increased fat mass composition, decreased lean mass composition, lower energy expenditure, and locomotor activity in both sexes. Male IGF1R/IR^LepRb^ mice exhibited impaired insulin sensitivity. These findings suggest that IGF1R and IR in LepRb neurons jointly regulated body composition, energy balance, and glucose homeostasis. Taken together, our studies identified the sex-dependent complex roles of IGF1R and IR in LepRb neurons in regulating body growth, reproduction, and metabolism.

## INTRODUCTION

Growth and reproduction are intricately linked (*1, 2*). Insulin-like growth factor 1 (IGF-1) is the major mediator of growth hormone (GH)-stimulated somatic growth, as well as GH-independent anabolic responses such as embryonic growth and reproductive function (*3*). IGF-1 administration advances pubertal timing (*4*). Notably, ablation of IGF-1 receptors (IGF1R) in the brain causes growth retardation, infertility, and glucose intolerance in mice (*5*). To narrow down the neurons that IGF1R acts through, a group of researchers deleted IGF1R in gonadotropin-releasing hormone (GnRH) neurons that control the maturation of the reproductive axis, and only found delayed puberty by 3-4 days with normal adult reproductive function (*4*). These findings suggested that other upstream neurons responsive to IGF-1 may alter GnRH neuronal activity and regulate reproductive function.

Neurons expressing the long form of the leptin receptor (LepRb) sense various metabolic cues to regulate various physical processes including puberty onset, adult fertility, energy balance, glucose homeostasis, and bone health (*6–18*). Many of the actions of leptin are attributable to effects in LepRb neurons, particularly in the mediobasal hypothalamus, including the arcuate nucleus (ARH) (*19, 20*). Disruption of ARH LepRb neurons causes modest weight gain (*21, 22*). LepRb neurons in the dorsomedial hypothalamus co-expressing *Glp1r* suppress food intake and body weight (*23*), and mediate leptin’s thermoregulatory actions (*24*). Unexpectedly, GH signaling in LepRb neurons did not influence body growth or food intake but played a critical role in regulating glucose metabolism (*25*). These findings led to our interest in studying the role of IGF1R in LepRb neurons in body growth, reproduction, and metabolism.

IGF-1 and insulin act through related tyrosine kinase receptors whose signals converge on downstream insulin receptor substrate (IRS) proteins (*26*) and then recruit and activate phosphatidylinositol 3-kinase (PI3K) to promote Akt signaling (*27*). Of the IRS-proteins, IRS2 pathways were found to integrate female reproduction and energy homeostasis, as mice lacking IRS2 displayed small, anovulatory ovaries with decreased numbers of follicles (*28*). Loss of IRS2 in LepRb neurons in mice led to obesity, glucose intolerance, and insulin resistance, but their reproductive capacity was normal (*12*). PI3K signaling in LepRb neurons plays an essential role in energy expenditure, reproduction, and body growth (*11*). Disruption of PI3K p110α and p110 β subunits increased energy expenditure, locomotor activity, and thermogenesis while delaying puberty and impairing fertility (*11*). Surprisingly, although deletion of IR in LepRb neurons caused a mild delay of puberty, it did not recapitulate the other metabolic and reproductive changes seen in PI3K knockout mice (*11*). IGF1R and IR compensate for each other to maintain normal muscle growth (*29*) and white and brown fat mass formation in mice (*30*). Therefore, we hypothesized that the IGF1R and IR in LepRb neurons jointly support metabolic and reproductive function. To test this hypothesis, we generated mice lacking IGF1R exclusively in LepRb neurons (IGF1R^LepRb^ mice) and mice simultaneously lacking both IGF1R and IR in LepRb neurons (IGF1R/IR ^LepRb^ mice), and then characterized the impact on the regulation of body growth, reproduction, and metabolism in these models.

## MATERIALS AND METHODS

### Animals and genotyping

To generate mice with the IGF1Rs specifically deleted in LepRb-expressing neurons, LepR-Cre mice (*31*) were crossed with IGF1R-floxed mice (*32, 33*) (RRID: IMSR_JAX:023426) and bred to homozygosity for the floxed allele only. The IGF1R^flox/flox^ mice were designed with loxP sites flanking exon 3. Excision of exon 3 in the presence of Cre recombinase results in a frame shift mutation and produces a premature stop codon. Littermates only carrying Cre recombinase were used as controls (LepR-Cre). To generate double-knockout of IGF1R and IR, LepR-Cre mice (*31*) were crossed with IGF1R-floxed and IR-floxed mice (*34*) and bred to homozygosity for the floxed alleles only. All mice were on a C57BL/6 background. Where specified, the mice also carried the reporter Ai14 (Jax line 007914) (*16, 35*), in which a loxP-flanked STOP cassette prevents transcription of a CAG promoter-driven red fluorescent protein (tdTomato) inserted into the ROSA26 locus. Mice were housed in the University of Toledo College of Medicine animal facility at 22°C to 24°C on a 12-hour light/12-hour dark cycle and were fed standard rodent chow (2016 Teklad Global 16% Protein Rodent Diet, 12% fat by calories; Harlan Laboratories, Indianapolis, Indiana). On postnatal day (PND) 21, mice were weaned. At the end of the study, all animals were sacrificed by CO_2_ asphyxiation or by cardiac puncture under 2% isoflurane anesthesia to draw blood. Mice were genotyped using the pairs of primers described in Table 1. The University of Toledo College of Medicine Institutional Animal Care and Use Committee approved all procedures.

**Table 1.**
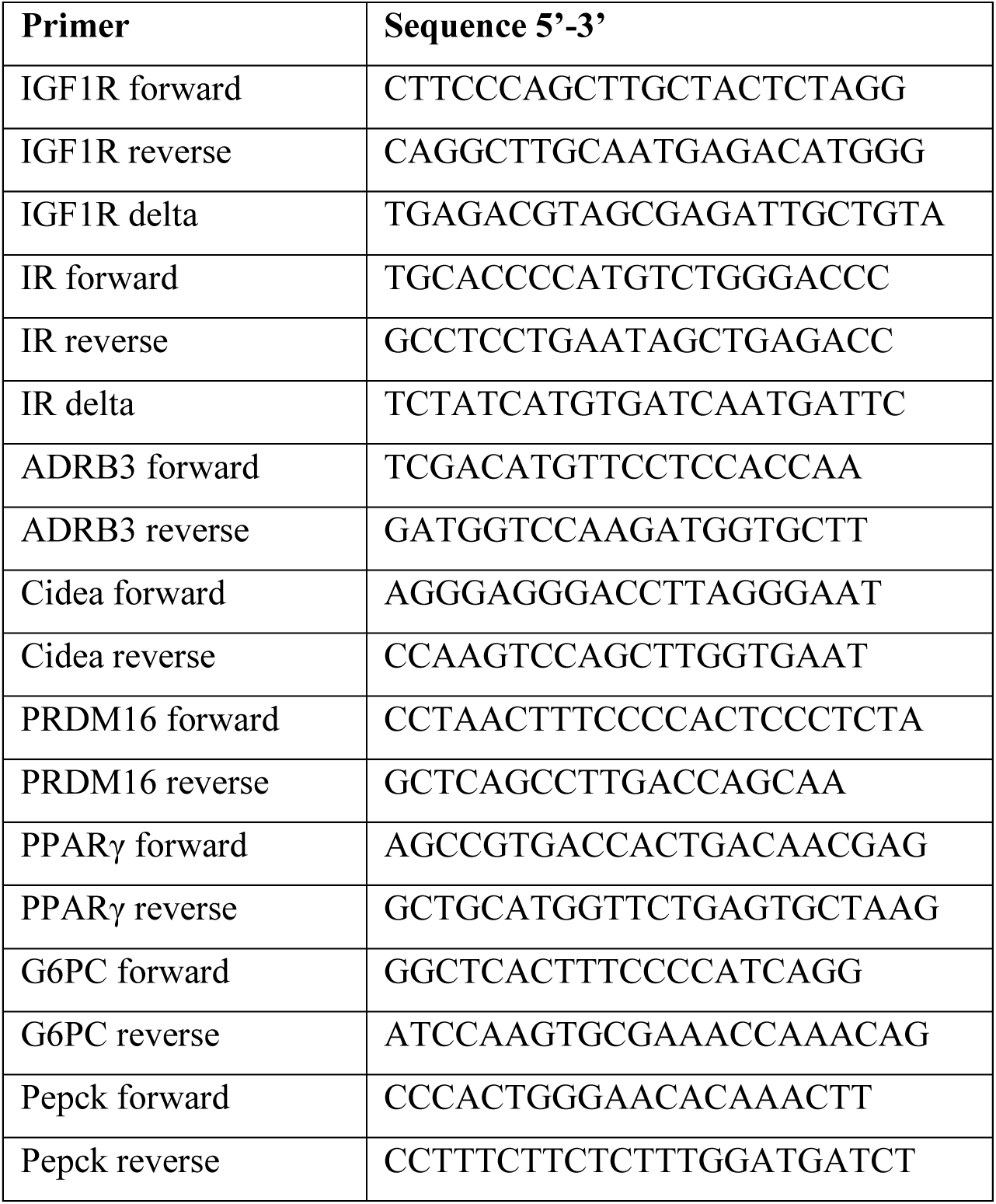
Summary of primers.

### Puberty and reproductive phenotype assessment

Timing of pubertal development was checked daily after weaning at 21 days by determining vaginal opening (VO) in female mice or balanopreputial separation (BPS) in male mice. Saline lavages were used to collect vaginal cells of female mice following VO. The first estrus age was identified as two consecutive days with keratinized cells after two previous days with leukocytes (*11*). Estrus stages were assessed based on vaginal cell cytology as described previously (*11, 36*). BPS was checked daily from weaning by manually retracting the prepuce with gentle pressure (*37*). After BPS was seen in male mice, each male mouse was paired with one fertile wild-type female to evaluate the first date of conception while monitoring daily for copulatory plugs. The paired mice were separated until males reached 8 weeks of age, and pregnancy rate, litter size, and interval from mating to birth were recorded. The age of sexual maturation was estimated from the birth of the first litter minus the average pregnancy duration for mice (21 days). At 3 months of age, we examined adult fertility. Animals were paired with fertile adult wild-type breeders for 8 nights to collect additional data on pregnancy rate, interval from mating to birth, and number of pups per litter. To examine the fertility at different ages, we examined the adult fertility at 4, 7, 10, 14, and 17 months of age; the pregnancy rate and number of pups per litter were recorded accordingly.

### Body length measurement

Body length, from the tip of the nose to the base of the tail was measured weekly from week 3 to 20 when mice were anesthetized under 2% isoflurane.

***Body composition assessment and indirect calorimetry***. Body weight was measured weekly from week 3 to 20. Body composition was assessed by nuclear magnetic resonance (Minispec mq7.5; Bruker Optics, Billerica, Massachusetts) to determine the percentage of fat mass, lean mass, and body fluid (*38*). We performed indirect calorimetry in mice at the age of 3 to 4 months in a Calorimetry Module (CLAMS; Columbus Instruments, Columbus, Ohio) as described previously (*39*). Adult (14- to 16-week-old) IGF1R^LepRb^, IGF1R/IR^LepRb,^ and age-matched control LepRb-Cre (n=7-11/genotype) males and females were weighed and then individually placed into the sealed chambers with free access to food and water. The study was conducted in an experimentation room set at 21°C-23°C with 12-hour-light/dark cycles. The metabolic assessments were conducted continuously for 72 hours after 24 hours of adaptation. The consumption of oxygen (VO_2_) and production of carbon dioxide (VCO_2)_ in each chamber was sampled sequentially for 1 minute in a 20-minute interval, and the motor activity was recorded every second in x and z dimensions. Respiratory exchange ratio was calculated as VCO_2_/VO_2_, and energy expenditure was calculated based on the formula: EE = 3.91 × [(VO_2_) + 1.1 × (VCO_2_)]/1000.

### Glucose tolerance test (GTT) and insulin tolerance test (ITT)

GTTs and ITTs were performed as described previously (*38*). For GTTs, after a 16-hour fast, mice were injected with dextrose (2g/kg i.p.). Tail blood glucose was measured using a veterinary glucometer (AlphaTRAK; Abbott Laboratories, Abbott Park, Illinois) before and 15, 30, 45, 60, 90, and 120 minutes after injection. For ITT, after a 3-hour fast, mice were injected with recombinant insulin (0.75 U/kg i.p.). Tail blood glucose was measured again at specified time points.

### Hormone assays

Submandibular blood was collected at 9:00 to 11:00 AM to detect basal luteinizing hormone (LH), follicle-stimulating hormone (FSH) and estradiol levels from mice between postnatal day 21 and 28 (before showing vaginal opening or balanopreputial separation) and 3-month-old male and female mice on diestrus. LH and FSH were measured via RIA performed by the University of Virginia Center for Research in Reproduction Ligand Assay and Analysis Core (Charlottesville, VA). The assay for LH had a detection sensitivity of 3.28 pg/ml. The intra-assay and inter-assay coefficients of variance (CVs) were 4.0% and 8.6%. The assay for FSH had a detection sensitivity of 7.62 pg/ml. The intra-assay and inter-assay CVs were 7.4% and 9.1%. Serum estradiol was measured by ELISA (Calbiotech, Spring Valley, California) with a sensitivity of 3 pg/mL and intra-assay and inter-assay CVs of <10%. Serum testosterone was measured by ELISA (Calbiotech, Spring Valley, California) with a sensitivity of 0.1 ng/mL and intra-assay and inter-assay CVs of <10%. Serum IGF-1 was measured by ELISA (Crystal Chem, Elk Grove Village, IL) with a sensitivity of 0.5 to 18 ng/mL and precision intra-assay and inter-assay CVs of <10%. Serum GH was measured by ELISA (Crystal Chem, Elk Grove Village, IL) with a sensitivity range of 0.15 to 9 ng/mL and intra-assay and inter-assay CVs of <10%. Serum insulin was measured by ELISA (Crystal Chem, Elk Grove Village, IL) with a sensitivity range of 0.1 to 12.8 ng/mL and intra-assay and inter-assay CVs of <10%. Serum C-Peptide was measured by ELISA (Crystal Chem, Elk Grove Village, IL) with a sensitivity range of 0.37 to 15 ng/mL and intra-assay and inter-assay CVs of <10%.

### Micro-computed tomography (mCT)

Dissected right femora and lumbar vertebrae from 5-month-old mice (n = 4/genotype) were immersed in 10% formalin and stored in the dark. To determine the tissue microarchitecture and densitometry, bones were scanned using the mCT-35 system (Scanco Medical AG, Bruettisellen, Switzerland), as previously described (*40*). Scan parameters included 7-micron nominal resolution with the X-ray source operating at 70 kVp, and a current of 113 µA. As described previously (*40*), scans of the proximal tibia consisted of 300 slices starting at the growth plate. Images of trabecular bone were segmented at 220 threshold values using per mille scale following manual contouring starting 10 slices below the growth plate and extending to the end of the image stack. Scans of cortical bone at the tibia midshaft consisted of 55 slices and images of cortical bone were contoured in the entire image stack and segmented at 260 thresholds using per mille scale. The analyses of the trabecular bone microstructure and the cortical bone parameters were performed using Evaluation Program V6.5-1 (Scanco Medical AG, Bruettisellen, Switzerland) and conformed to recommended guidelines (*41*). All mCT measurements were performed in a blind fashion.

### Tissue collection and histology

Ovaries, testes, white adipose tissue, and brown adipose tissue were collected from mice and fixed immediately in 10% formalin overnight and then transferred to 70% ethanol. Then tissues were embedded in paraffin and cut into 5- to 8-µm sections. Sections were stained by hematoxylin and eosin and then analyzed. For follicle and sperm quantification, a minimum of four ovaries and testes for each genotype at 5-month-old age were collected. Follicles were classified into the following categories: primordial, primary, secondary, and Graafian. Testes sections were analyzed by evaluating sperm stages, including counting the number of spermatogonium, spermatocytes, spermatid, and spermatozoa using light microscopy under 20x magnification (*42*). Sperm counts are reported per seminiferous tubule cross-section.

### Quantitative real-time PCR

Mice were placed under isoflurane anesthesia followed by decapitation and removal of the hypothalamus. Total hypothalamic RNA was extracted from dissected tissues by an RNeasy Lipid Tissue Mini Kit (QIAGEN, Valencia, California) (*43*). Single-strand cDNA was synthesized by a High-Capacity cDNA Reverse Transcription kit (Applied Biosystems) using random hexamers as primers as listed in Appendix A. Each sample was analyzed in duplicate to measure gene expression level. A 25 µM cDNA template was used in a 25 µl system in 96-well plates with SYBR Green qPCR SuperMix/ROX (Smart Bioscience Inc, Maumee, Ohio). The reactions were run in an ABI PRISM 7000 sequence detection system (PE Applied Biosystems, Foster City, California), or a 10 µM cDNA template was used in a 10 µl system in 384-well plates with SYBR Green qPCR SuperMix/ROX (Smart Bioscience Inc, Maumee, Ohio). These reactions were run in a ThermoFisher QuantStudio 5 Real-Time PCR system (Applied Biosystems, Foster City, California). All data were analyzed using the comparative Ct method (2^-ΔΔCt^) with glyceraldehyde-3-phosphate dehydrogenase (GADPH) as the housekeeping gene. The mRNA expression in IGF1R^LepRb^ and IGF1R/IR^LepRb^ versus LepRb-Cre control mice was determined by a comparative cycle threshold method and relative gene copy number was calculated as 2^-ΔΔCt^ and presented as fold change from the relative mRNA expression of the LepRb-Cre control group.

### Perfusion and immunohistochemistry

Adult LepRb-Cre, IGF1R^LepRb,^ and IGF1R/IR^LepRb^ mice at the ages of 3 to 6 months were deeply anesthetized by ketamine and xylazine. After brief perfusion with a saline rinse, mice were perfused transcardially with 10% formalin for 10 minutes, and the brain was removed. The brain was postfixed in 10% formalin at 4°C overnight and immersed in 10%, 20%, and 30% sucrose at 4°C for 24 hours each. Then 30-µm sections were cut by a sliding microtome into five equal serial sections. After rinsing in PBS, sections were blocked for 2 hours in PBS-T (PBS, Triton X-100, and 10% normal horse serum). Samples were incubated for 48 hours at 4°C in PBS-T-containing rabbit anti-IGF1R β antibody (1:1000; Cell signaling, Cat#9750). After three washes in PBS, sections were incubated in PBS-T (Triton X-100 and 10% horse serum) containing secondary antibody Alexa Flour 488 (1:1,000, Thermofisher Scientific, Cat. #A-21206) for 2 hours at room temperature. Finally, sections were washed, mounted on slides, cleared, and coverslipped with fluorescence mounting medium containing DAPI (Vectasheild, Vector Laboratories, Inc. Burlingame, California).

### Statistical analysis

Data are presented as mean ± SEM. Normality testing was used to determine the normal distribution of data. If the data followed a normal distribution, One-way ANOVA was used as the main statistical method to compare the three groups, followed by the Tukey multiple comparison test.

If the data did not follow a normal distribution, the Kruskkal-Wallis test was used. For body weight, body length, GTTs, and ITTs, a two-way ANOVA was used to compare changes over time among three groups. Bonferroni multiple comparison tests were then performed to compare differences between groups. A value of *P* ≤ 05 was considered to be significant.

## RESULTS

### Disruption of IGF1R in IGF1R^LepRb^ mice

Genetic ablation of IGF1R in LepRb neurons was validated using immunostaining (Fig. 1A). Approximately 15% of LepRb neurons express IGF1R protein. Compared to control mice, colocalization was sharply lower in IGF1R^LepRb^ mice (Fig. 1B). Hypothalamic IGF1R mRNA expression was lower in IGF1R^LepRb^ mice and IGF1R/IR^LepRb^ mice (Fig. 1C). Hypothalamic IR mRNA expression was lower in IGF1R/IR^LepRb^ mice (Fig. 1D). No changes were seen in GHR mRNA expression (Fig. 1E).

**Figure 1.**
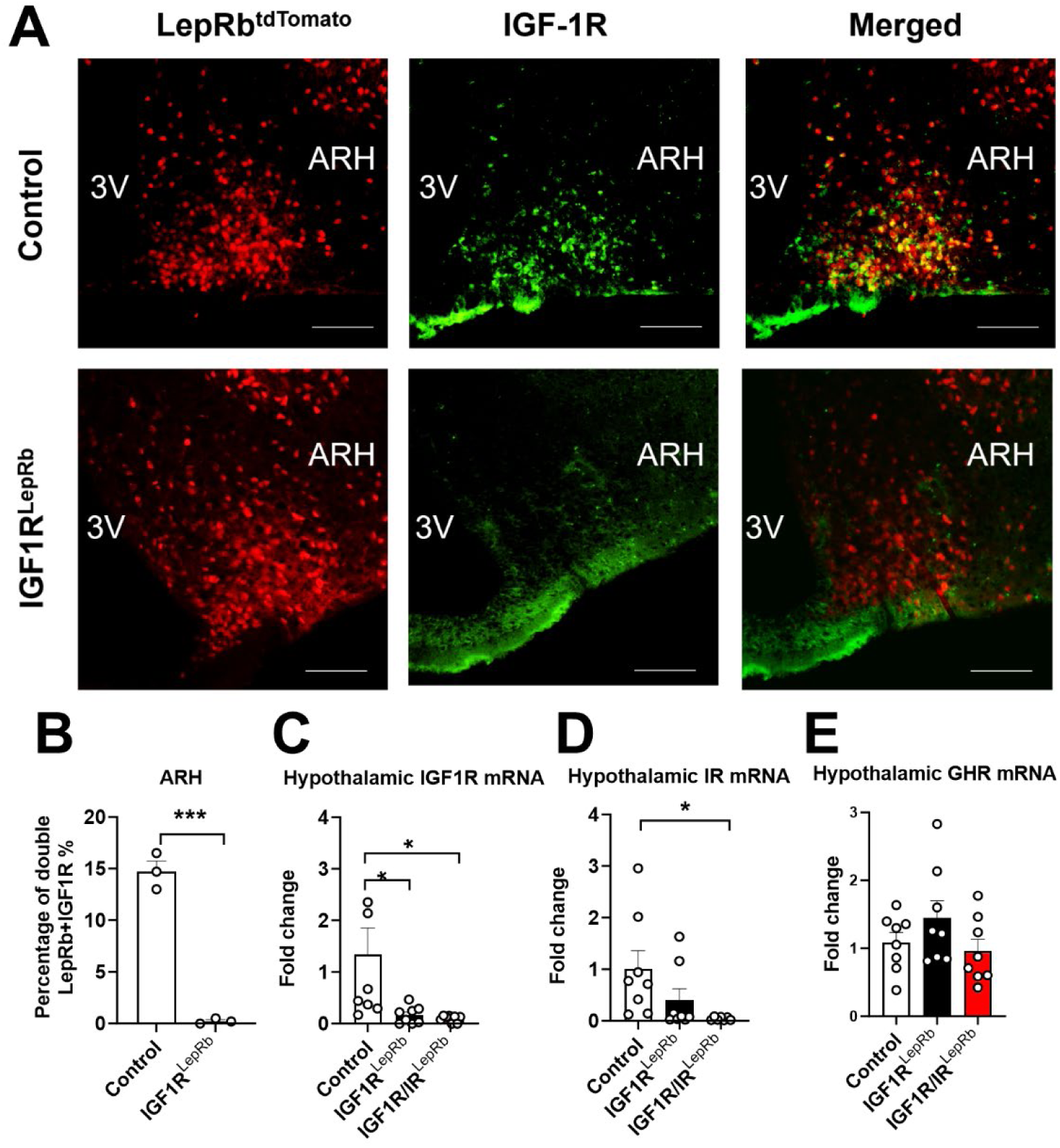
Confirmation of mouse model. (A) Colocalization of LepRb neuron and IGF1Rs from control and IGF1R^LepRb^ mice. (B) Quantification of colocalization (n=3/group). (C-E) Hypothalamic IGF1R (C), IR (D), and GHR (E) mRNA expression were evaluated in control, IGF1RLepRb, and IGF1R/IRLepRb mice (n=8/group). Values throughout figure are means ±SEM. For the entire figure, \**P* < 0.05, ** *P* < 0.01, and *** *P* < 0.0001, were determined by Tukey’s post hoc test following one-way ANOVA for each group.

### Delayed Puberty and impaired fertility in IGF1R^LepRb^ and IGF1R/IR^LepRb^ mice

Female IGF1R^LepRb^ mice had significantly delayed vaginal opening age (at 30.8±0.5 days in IGF1R^LepRb^ vs 37.0±1.0 days in controls) and first estrus age (at 37.6±0.4 days in IGF1R^LepRb^ vs 42.1±0.5 days in controls) (Fig. 2A-B). No changes were seen in estrus cycle length or time spent in each estrous stage (Fig. 2C-D). Female IGF1R^LepRb^ mice at 3 months of age had significantly impaired fertility and lower numbers of pups per litter (Fig. 2E-F). We did not find alterations in serum levels of LH, FSH, or estradiol (Fig. 2G-I). No difference was seen in ovary weight (Fig. 2J). However, we found a lower number of Graafian follicles in 5-month-old female IGF1R^LepRb^ mice (Fig. 2K-N), which may be related to reproductive deficits. These findings indicate that IGF1R signaling in LepRb neurons is indispensable for normal puberty onset and adulthood fertility in female mice. The reproductive phenotype seen in female IGF1R/IR^LepRb^ mice was comparable to IGF1R^LepRb^ mice except for the significantly later first estrus age (Fig. 2A-K), suggesting IR signaling in LepRb neurons also regulates the onset of puberty.

**Figure 2.**
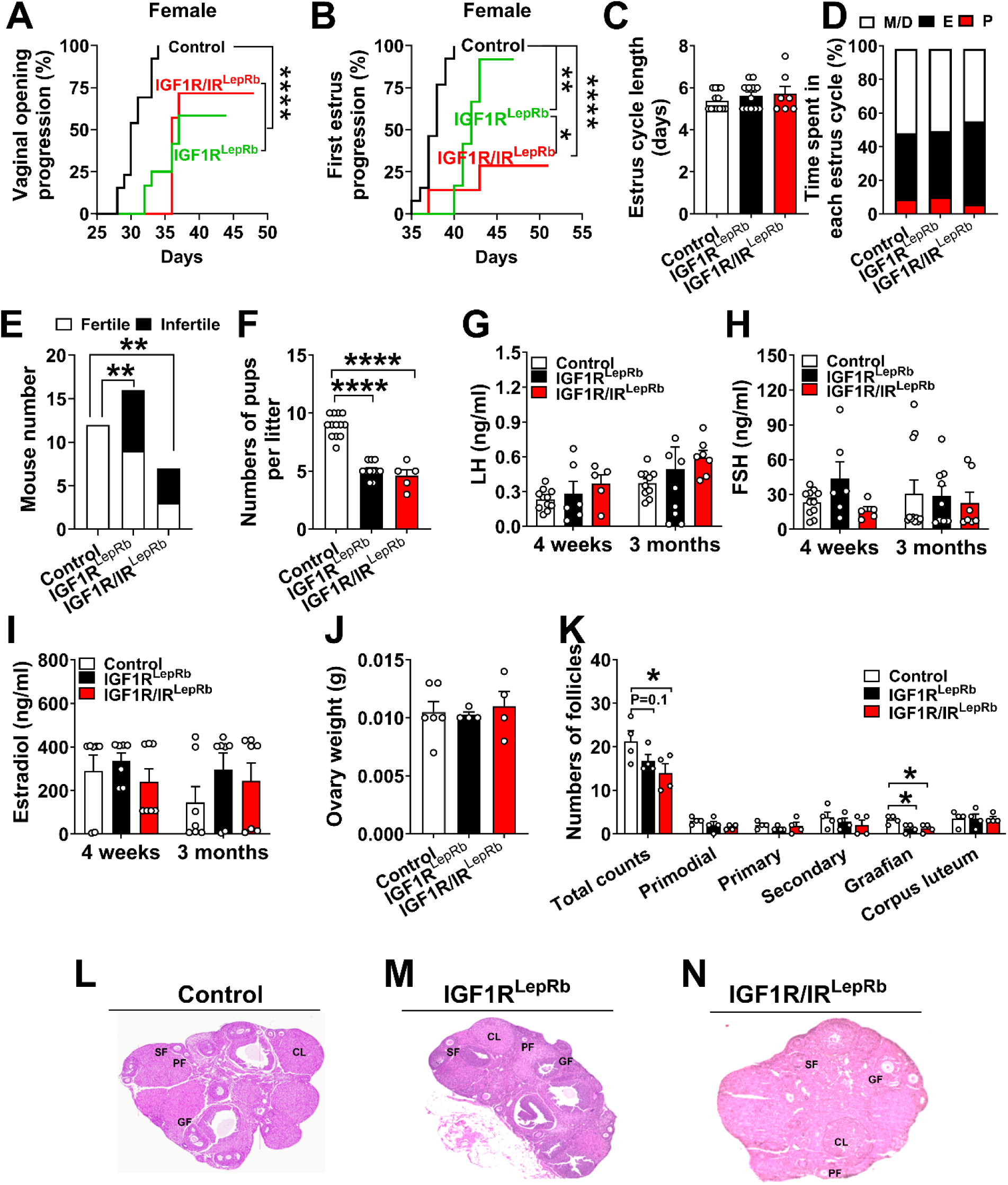
Reproductive deficits in female IGF1R^LepRb^ and IGF1R/IR^LepRb^ mice. (A) Vaginal opening age, (B) first estrus age, (C) estrus cycle length, and (D) estrus cyclicity were evaluated in female control, IGF1R^LepRb,^ and IGF1R/IR^LepRb^ mice (n=7-13/group). (E) Pregnancy rate and (F) number of pups per litter in 4-month-old female control, IGF1R^LepRb^ and IGF1R/IR^LepRb^ mice (n=7-13/group). (G) Serum LH, (H) FSH, and (I) estradiol levels on diestrus in 4-week-old and 3-month-old female control, IGF1R^LepRb^ and IGF1R/IR^LepRb^ mice (n=5-11/group). (J) Ovary weight, (K) histological analysis of ovarian follicles (n=4-6/group) and (L-N) representative sliced and HE-stained paraffin-embedded ovaries in 5-month-old female control, IGF1R^LepRb^ and IGF1R/IR^LepRb^ mice. PF, primary follicle; SF, secondary follicle; GF, Graafian follicle; CL, corpus luteum. Values throughout the figure are means ±SEM. For the entire figure, \**P* < 0.05, ** *P* < 0.01, *** *P* < 0.0001, **** *P* < 0.00001, determined by Log Rank Test or Chi-square or Tukey’s post hoc test following one-way ANOVA.

Male IGF1R^LepRb^ mice had significantly delayed balanopreputial separation age (33.8±0.9 days of age in IGF1R^LepRb^ vs 41.5±0.6 days in controls) and first date of conception (49.8±0.9 days of age in IGF1R^LepRb^ vs 57.4±1.1 days in controls) (Fig. 3A-B). At 3 months of age, male IGF1R^LepRb^ mice had significantly impaired fertility and lower numbers of pups per litter (Fig. 3C-D). In addition, loss of IGF1R signaling in LepRb neurons also caused significantly lower levels of LH and FSH at 3 months and lower testosterone levels at 4 weeks in male mice (Fig. 3E-G). The testis histological analysis showed a significantly decreased number of spermatids and spermatozoa in the seminiferous tubules of 5-month-old male IGF1R^LepRb^ mice (Fig. 3H). Figure 3I-K showed representative images of seminiferous tubules. Thus, like female IGF1R^LepRb^ mice, IGF1R signaling in LepRb neurons also plays a dominant role in the regulation of reproductive function in male mice. Male IGF1R/IR^LepRb^ mice showed more profound reproductive deficits (lower number of pups per litter and lower FSH levels) than IGF1R^LepRb^ mice (Fig. 3D and H). Together, these results suggest that both IGF1R and IR signaling in LepRb neurons are required for normal reproductive function in female and male mice.

**Figure 3.**
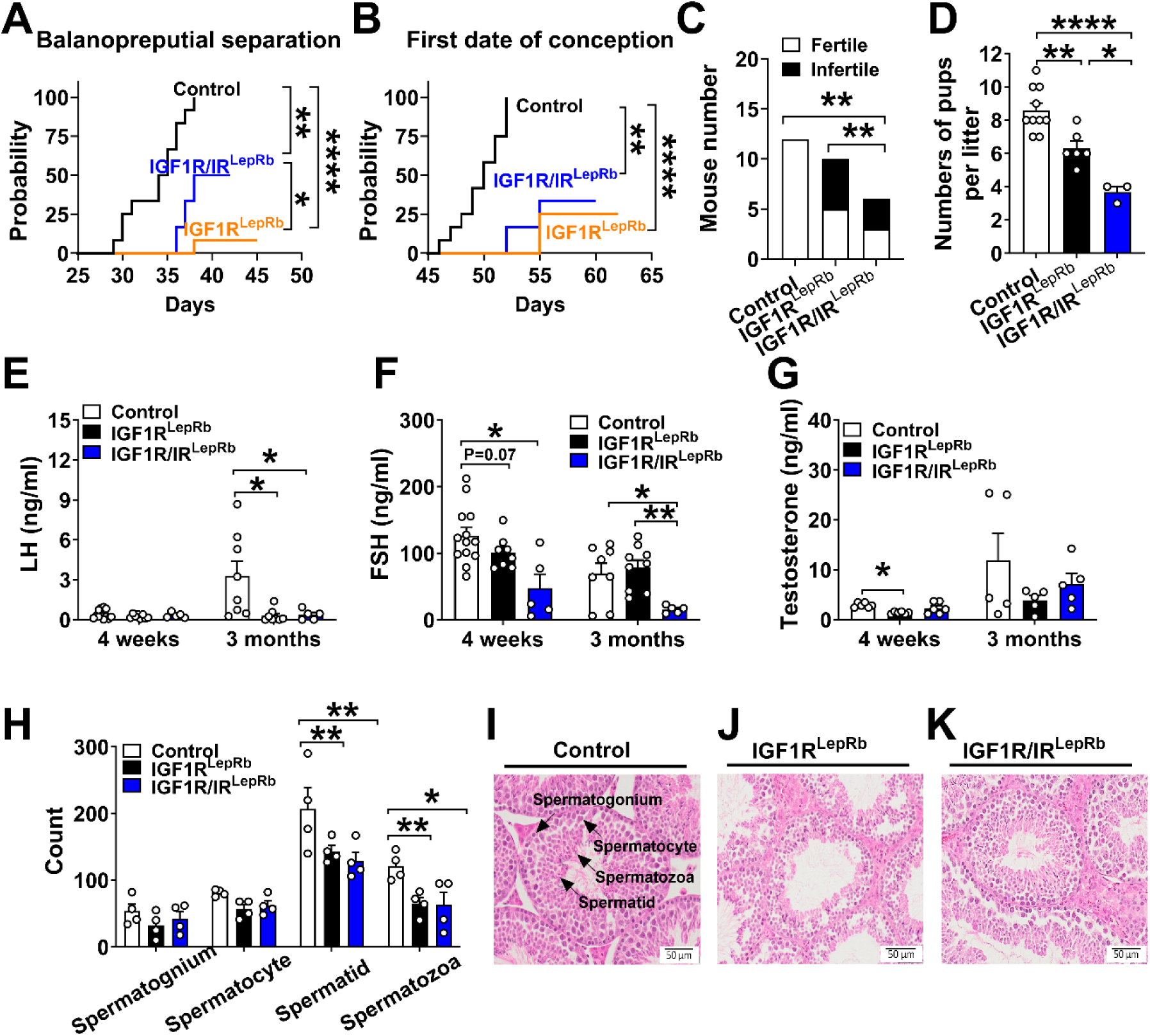
Reproductive deficits in male IGF1R^LepRb^ and IGF1R/IR^LepRb^ mice. (A) Balanopreputial separation age and (B) first date of conception in male control, IGF1R^LepRb^, and IGF1R/IR^LepRb^ mice (n=6-12/group). (C) Pregnancy rate and (D) numbers of pups per litter in male control, IGF1R^LepRb^, IGF1R/IR^LepRb^ mice (n=6-12/group). (E) Serum LH, (F) FSH, and (G) testosterone levels in 4 weeks-old and 3 months-old male control, IGF1R^LepRb^ and IGF1R/IR^LepRb^ mice (n=5-12/group). (H) Analysis of cross- sectional testes seminiferous tubule sperm stages (n=4/group) and (I-K) representative sliced and HE-stained paraffin-embedded testes in 5-month-old male control, IGF1R^LepRb^ and IGF1R/IR^LepRb^ mice. Values throughout figure are means ±SEM. For the entire figure, \**P* < 0.05, ** *P* < 0.01, and *** *P* < 0.0001, were determined by Log Rank Test or Chi-square or Tukey’s post hoc test following one-way ANOVA.

A recent study reported that IGF-1 gene therapy induces GnRH release in the median eminence and maintains kisspeptin production in middle-aged female rats (*44*), suggesting IGF-1 may have a protective effect against reproductive decline. To explore the role of IGF1R signaling in LepRb neurons in the reproductive aging process, we performed fertility tests at 4, 7, 10, 14, and 17 months in IGF1R^LepRb^ mice and control mice. At 4-month-old age, female IGF1R^LepRb^ mice had impaired fertility and lower numbers of pups per litter (Fig. 4A and B), which was consistent with previous findings. The fertility in female IGF1R^LepRb^ mice declined to zero at 10 months of age while controls still maintained a nearly 50% pregnancy rate (Fig. 4A). Similarly, from 10 months onwards, the rate of fertility decline was faster in male IGF1R^LepRb^ mice compared to controls (Fig. 4C-D). Together, these results suggest that IGF1R in LepRb neurons might have a protective effect against reproductive aging.

**Figure 4.**
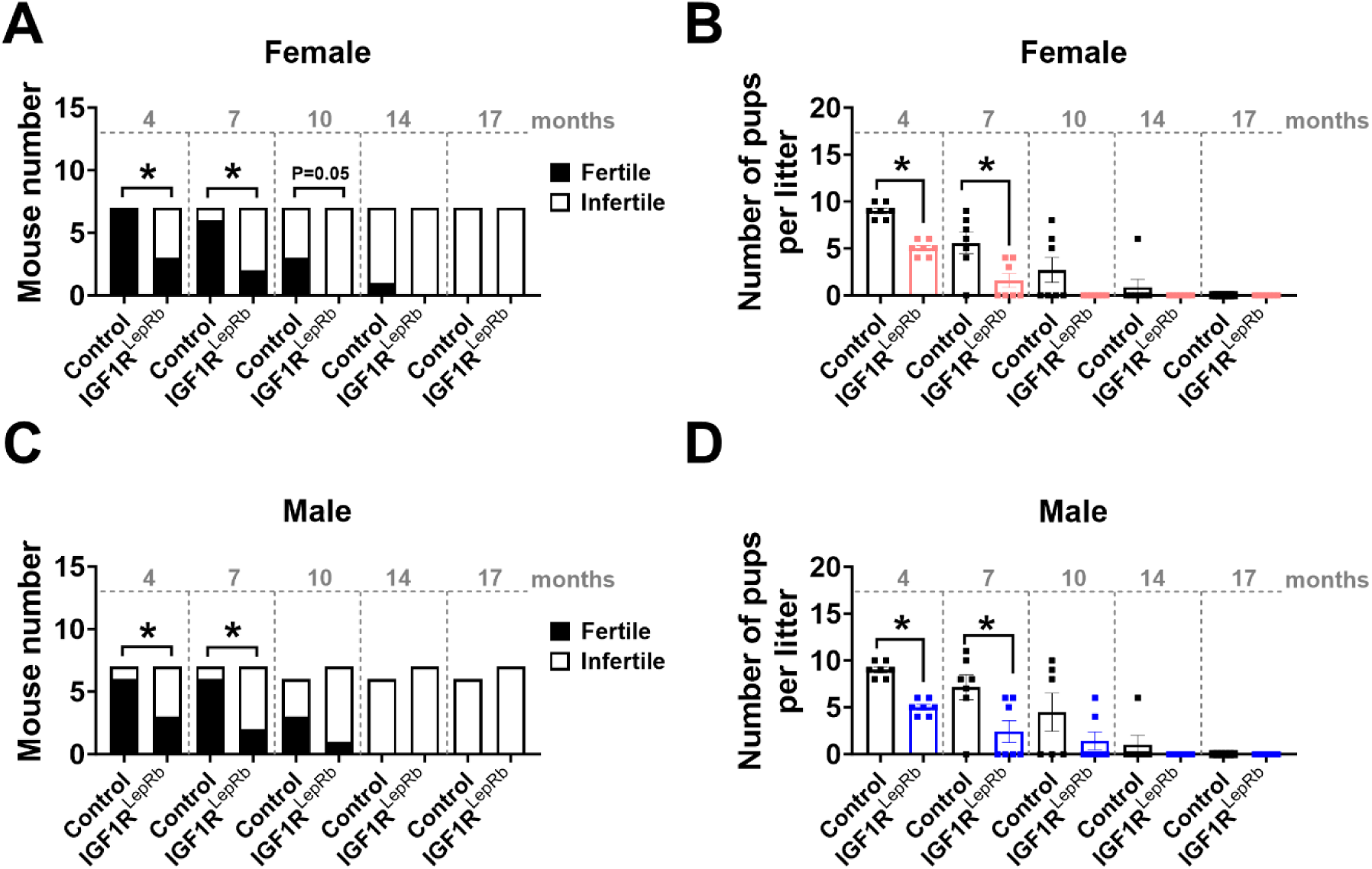
Advanced reproductive aging in IGF1R^LepRb^ mice. (A) Pregnancy rate and (B) litter size were measured in 4-, 7 -, 10-, 14- and 17-month-old female control and IGF1R^LepRb^ mice (n=7/group). (C) Pregnancy rate and (D) litter size in 4-, 7 -, 10-, 14- and 17-month-old male control and IGF1R^LepRb^ mice (n=7/group). Values throughout figure are means ±SEM. For the entire figure, \**P* < 0.05, determined by multiple t test.

### Body growth in IGF1R^LepRb^ and IGF1R/IR^LepRb^ mice and bone phenotype of IGF1R^LepRb^ mice

To determine whether IGF1R signaling in LepRb neurons regulates body growth and bone health, we measured body length and serum biomarkers and performed X-ray micro-computed tomography (mCT). Female IGF1R^LepRb^ mice had temporary growth retardation during weeks 3 to 6 (Fig. 5A), but no alterations in serum levels of IGF-1 or GH at the ages of 4 weeks and 3 months old (Fig. 5B-C). Female IGF1R/IR^LepRb^ mice displayed more profound but still temporary growth retardation compared to IGF1R^LepRb^ mice, with no changes in serum IGF-1 or GH (Fig. 5A-C). Male IGF1R^LepRb^ and IGF1R/IR^LepRb^ mice displayed temporary growth retardation (Fig. 5D), which was associated with decreased levels of IGF-1 and GH at 3-month-old age (Fig. 5E-F). These findings suggest that IGF1R signaling in LepRb neurons is critical for normal body growth and hormonal regulation in males. The additional loss of IR signaling in LepRb neurons did not induce robust changes in male mice compared to IGF1R^LepRb^ mice, further suggesting that the role of IGF1R signaling in LepRb neurons in body growth and hormonal regulation is unique.

**Figure 5.**
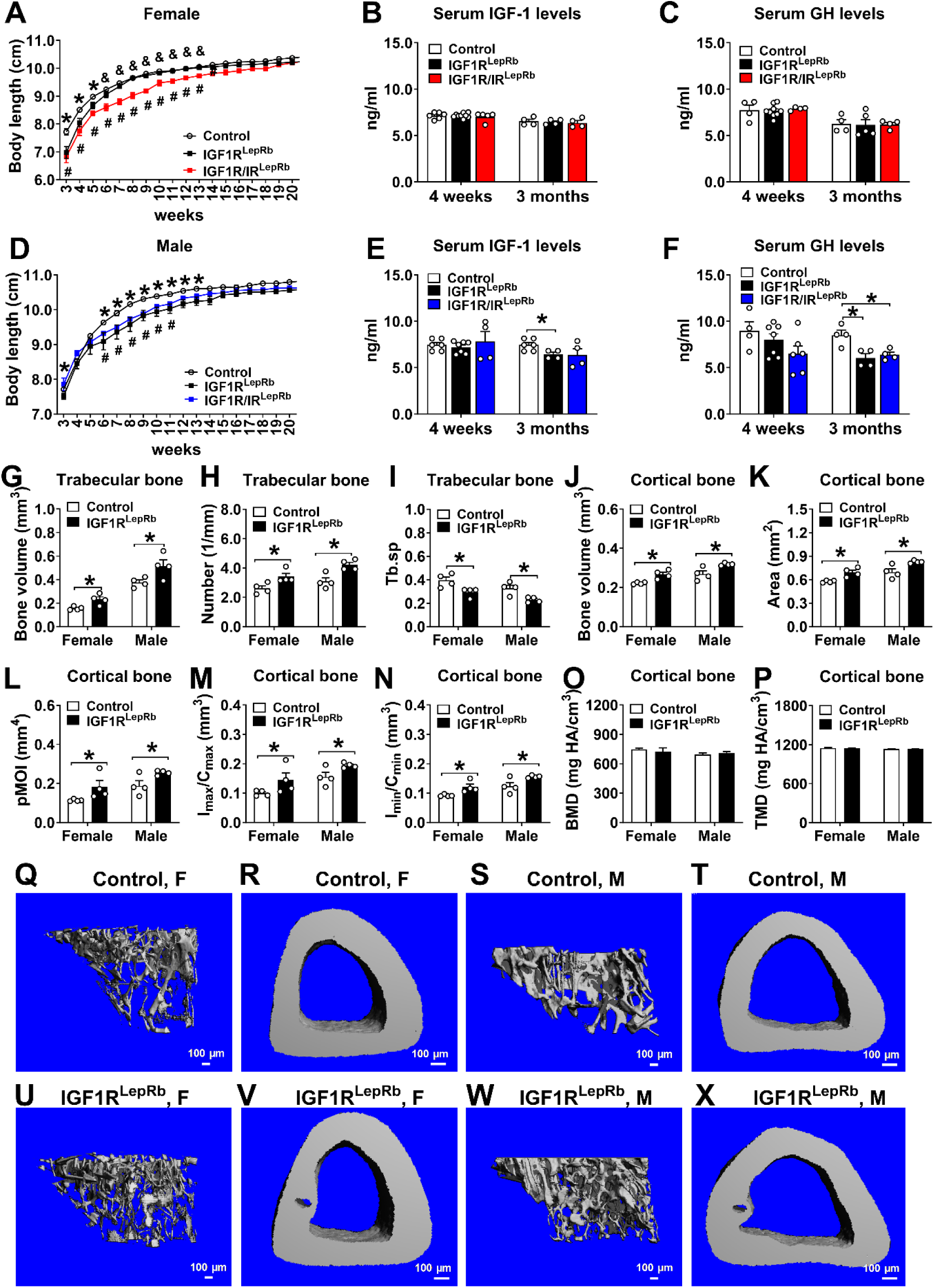
Body growth and bone phenotype in mice. (A) Body length curves from week 3 to 20 in female control, IGF1R^LepRb^, IGF1R/IR^LepRb^ mice (n=7-13/group). (B) Serum levels of IGF-1 and (C) GH at 4 weeks and 3 months of age in female control female control, IGF1R^LepRb^, and IGF1R/IR^LepRb^ mice (n=4-9/group). (D) Body length curves from week 3 to 20 in male control, IGF1R^LepRb^, IGF1R/IR^LepRb^ mice (n=7-13/group). (E) Serum levels of IGF-1 and (F) GH at 4 weeks and 3 months of age in male control female control, IGF1R^LepRb^, and IGF1R/IR^LepRb^ mice (n=4-7/group). (G) Trabecular bone volume, (H) numbers (Tb.N), (I) spacing (Tb.sp) and (J) cortical bone volume, (K) area (B.Ar), (L) polar moment of inertia (pMOI), (M) resistance to bending across the maximal (I_max_/C_max_), (N) minimal centroidedge (I_min_/C_min_), (O) Bone mineral density (BMD) and (P) tissue mineral density (TMD) in female and male control and IGF1R^LepRb^ mice (n=4/group). (Q-X) Representative images of trabecular and cortical bone in female and male control and IGF1R^LepRb^ mice. Values throughout figure are means ±SEM. For the entire figure, \**P* < 0.05 (control vs IGF1R^LepRb^ mice), ^#^*P* < 0.05 (control vs IGF1R/IR^LepRb^ mice), ^&^*P* < 0.05 (IGF1R^LepRb^ vs IGF1R/IR^LepRb^ mice), determined by t-test, Tukey’s post hoc test following one-way ANOVA or Bonferroni’s multiple comparison test following two-way ANOVA.

We next examined the bone phenotype using mCT in IGF1R^LepRb^ mice of 5 months of age. Compared to controls, female and male IGF1R^LepRb^ mice both displayed significantly increased trabecular bone volume, number, and spacing (Tb.Sp) (Fig. 5G-I) and cortical bone volume, area, polar moment of inertia (pMOI), resistance to bending across the maximal (I_max_/C_max_), and minimal centroidedge (I_min_/C_min_) (Fig. 5G-N) (Interestingly, there is no difference between sexes. For discussion, others showed sexual divergence in LepRb signaling in bone (*45*). However, no changes in bone mineral density (BMD) or tissue mineral density (TMD) were seen in IGF1R^LepRb^ mice of both sexes (Fig. 5 O-P). Figure 5Q-X shows representative renderings of trabecular bone and cortical bone in female and male control and IGF1R^LepRb^ mice. Together, our results suggest that IGF1R in LepRb expressing cells regulates body growth and bone mass accrual in mice. Unfortunately, no bone analysis was performed in IGF1R/IR^LepRb^ mice.

### Assessment of energy homeostasis in IGF1R^LepRb^ and IGF1R/IR^LepRb^ mice

We also evaluated the metabolic function of IGF1R^LepRb^ and IGF1R/IR^LepRb^ mice. Female IGF1R^LepRb^ mice had mildly decreased body weights and no changes in body composition (Fig. 6A-C). Female IGF1R^LepRb^ mice showed significantly decreased food intake with comparable energy expenditure but significantly increased locomotor activity (Fig. 6D-H). This phenotype was associated with elevated mRNA expression of adrenoceptor Beta 3 (ADRB3), cell death activator (Cidea), and PR-domain containing 16 (PRDM16) in brown adipose tissue (BAT) (Fig. 6I). No changes of weight, numbers of droplets, droplet area or histology of BAT were seen (Fig. 6J-M). Female IGF1R/IR^LepRb^ mice only showed temporarily lower body weight than IGF1R^LepRb^ mice (Fig. 6A). Interestingly, female IGF1R/IR^LepRb^ mice had dramatically increased percentage and absolute values of fat mass and decreased percentage and absolute values of lean mass (Fig. 6B-C). These mice also had significantly lower energy expenditure and locomotor activity (Fig. 6D-H), associated with decreased BAT weight and increased lipid droplet area in BAT (Fig. 6J-M). These results indicate that IGF1R and IR in LepRb neurons jointly regulate body composition, energy expenditure, and whitening of BAT.

**Figure 6.**
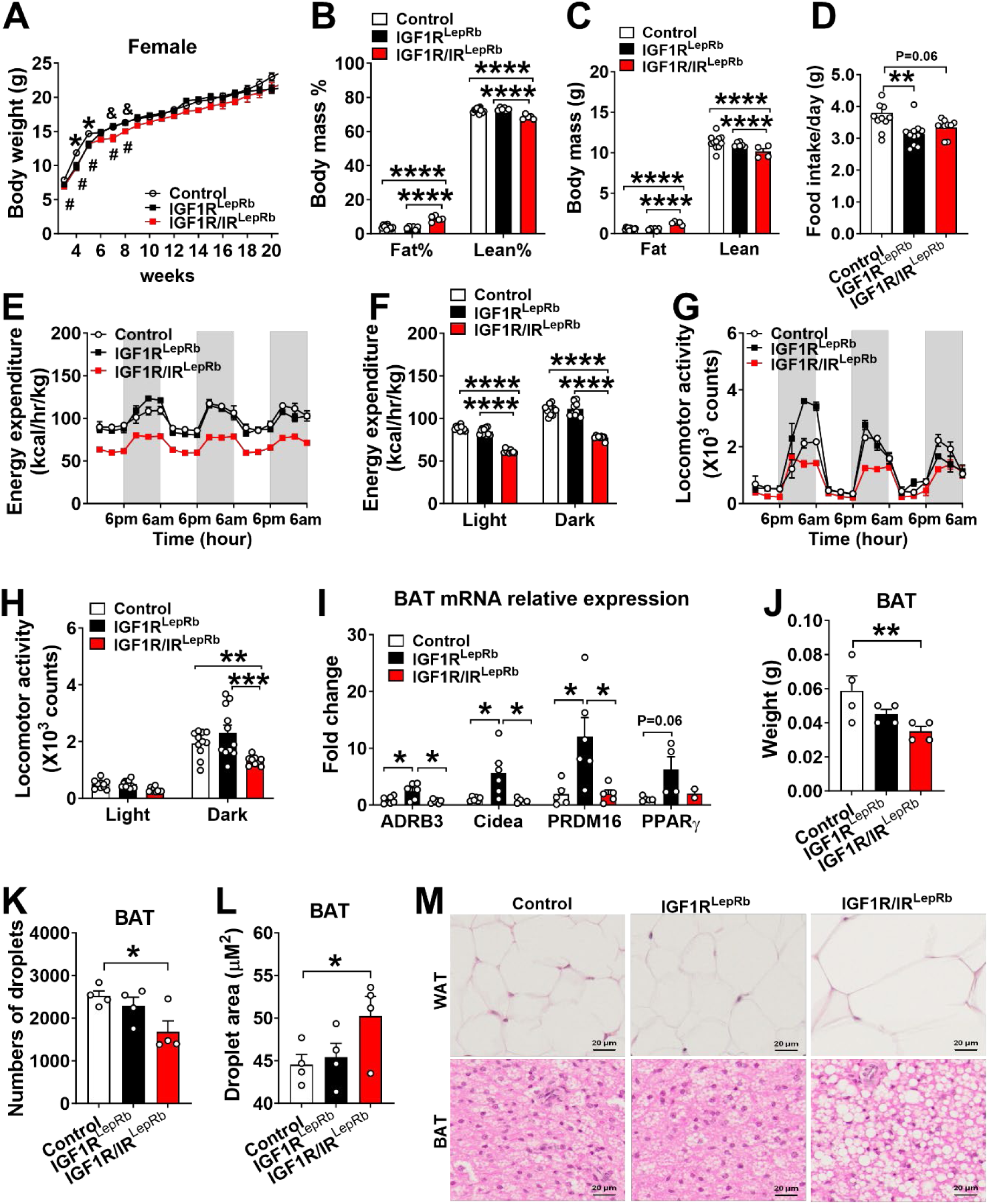
Altered energy homeostasis in female IGF1R^LepRb^ and IGF1R/IR^LepRb^ mice. (A) Body weight curves from week 3 to 20 in female control, IGF1R^LepRb,^ and IGF1R/IR^LepRb^ mice (n=7-13/group). (B) Body fat mass percentage, (C) lean mass percentage, and (D) food intake in 2-month-old female control, IGF1R^LepRb^ and IGF1R/IR^LepRb^ mice (n=5-11/group). (E-F) Energy expenditure and (G-H) physical activity in 3-month-old female control, IGF1R^LepRb^ and IGF1R/IR^LepRb^ mice (n=9-11/group). (I) Relative expression of thermogenesis markers as measured by quantitative PCR in brown adipose tissue (BAT) and (J) BAT weight in 5-month-old female control, IGF1R^LepRb,^ and IGF1R/IR^LepRb^ mice (n=4-6/group). (K) Numbers of droplets and (L) droplet area in BAT (n=4/group) and sliced and HE-stained paraffin-embedded BAT (M-O) in 5-month-old female control, IGF1R^LepRb^ and IGF1R/IR^LepRb^ mice. Values throughout figure are means ±SEM. For the entire figure, \**P* < 0.05, ** *P* < 0.01, *** *P* < 0.0001, and \*\*\*\**P* < 0.00001, were determined by Tukey’s post hoc test following one-way ANOVA or Bonferroni’s multiple comparison test following two-way ANOVA.

In contrast to females, male IGF1R^LepRb^ mice had no changes in body weight, body composition, food intake, energy expenditure, or thermogenic gene expression. Nevertheless, a decrease in locomotor activity was seen (Fig. 7A-J). Like female IGF1R/IR^LepRb^ mice, male IGF1R/IR^LepRb^ mice also showed dramatically increased fat mass percentage and decreased lean mass percentage and energy expenditure (Fig. 7B-F).

**Figure 7.**
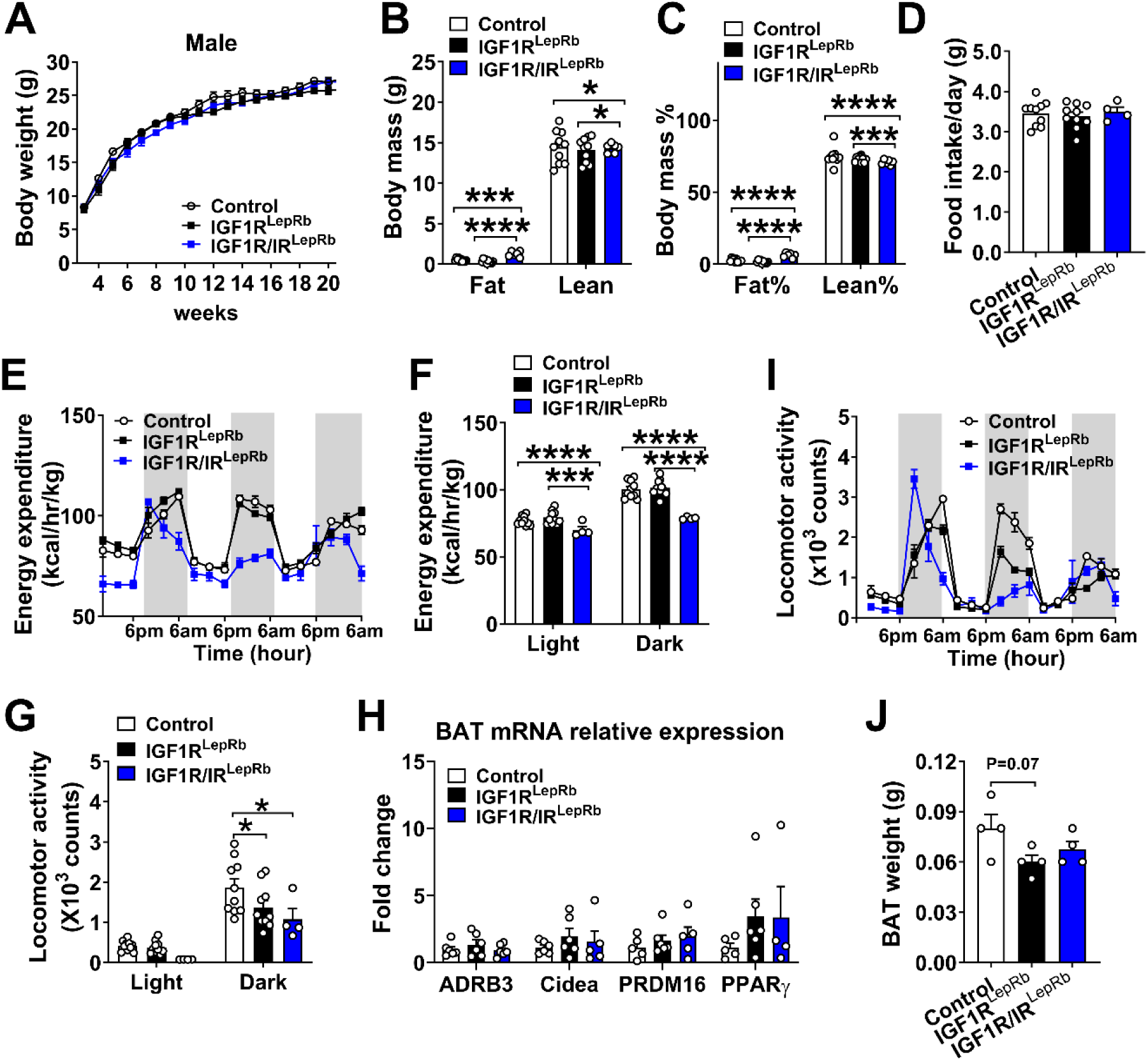
Altered energy homeostasis in male IGF1R^LepRb^ and IGF1R/IR^LepRb^ mice. (A) Body weight curves from week 3 to 20 in male control, IGF1R^LepRb,^ and IGF1R/IR^LepRb^ mice (n=6-10/group). (B) Body fat mass percentage, (C) lean mass percentage, and (D) food intake in 2-month-old male control, IGF1R^LepRb^ and IGF1R/IR^LepRb^ mice (n=4-10/group). (EF) Energy expenditure and (G-H) physical activity in 3-month-old male control, IGF1R^LepRb^ and IGF1R/IR^LepRb^ mice (n=4-10/group). (I) Relative expression of thermogenesis markers as measured by quantitative PCR in brown adipose tissue (BAT) and (J) BAT weight in 5-month-old male control, IGF1R^LepRb^ and IGF1R/IR^LepRb^ mice (n=4-6/group). Values throughout figure are means ±SEM. For the entire figure, \**P* < 0.05, ** *P* < 0.01, *** *P* < 0.0001, and \*\*\*\**P* < 0.00001, were determined by Tukey’s post hoc test following one-way ANOVA or Bonferroni’s multiple comparison test following two-way ANOVA.

### Assessment of glucose homeostasis in IGF1R^LepRb^ and IGF1R/IR^LepRb^ mice

To determine whether loss of IGF1R and/or IR in LepRb neurons causes an increased risk of diabetes, we evaluated glucose homeostasis in IGF1R^LepRb^ and IGF1R/IR^LepRb^ mice. Female IGF1R^LepRb^ mice showed glucose intolerance at 30 min and 45 min during the GTT but the area under curve (AUC) was not significantly changed (Fig. 8A-B). No changes were seen in ITT (Fig. 8C-D). Serum levels of insulin (Supplemental Fig. 1A), C-peptide (Supplemental Fig. 1B), insulin/C-peptide ratio (Fig. 8E), and insulin sensitivity as calculated by the homeostatic model assessment for insulin resistance (HOMA-IR)

**Figure 8.**
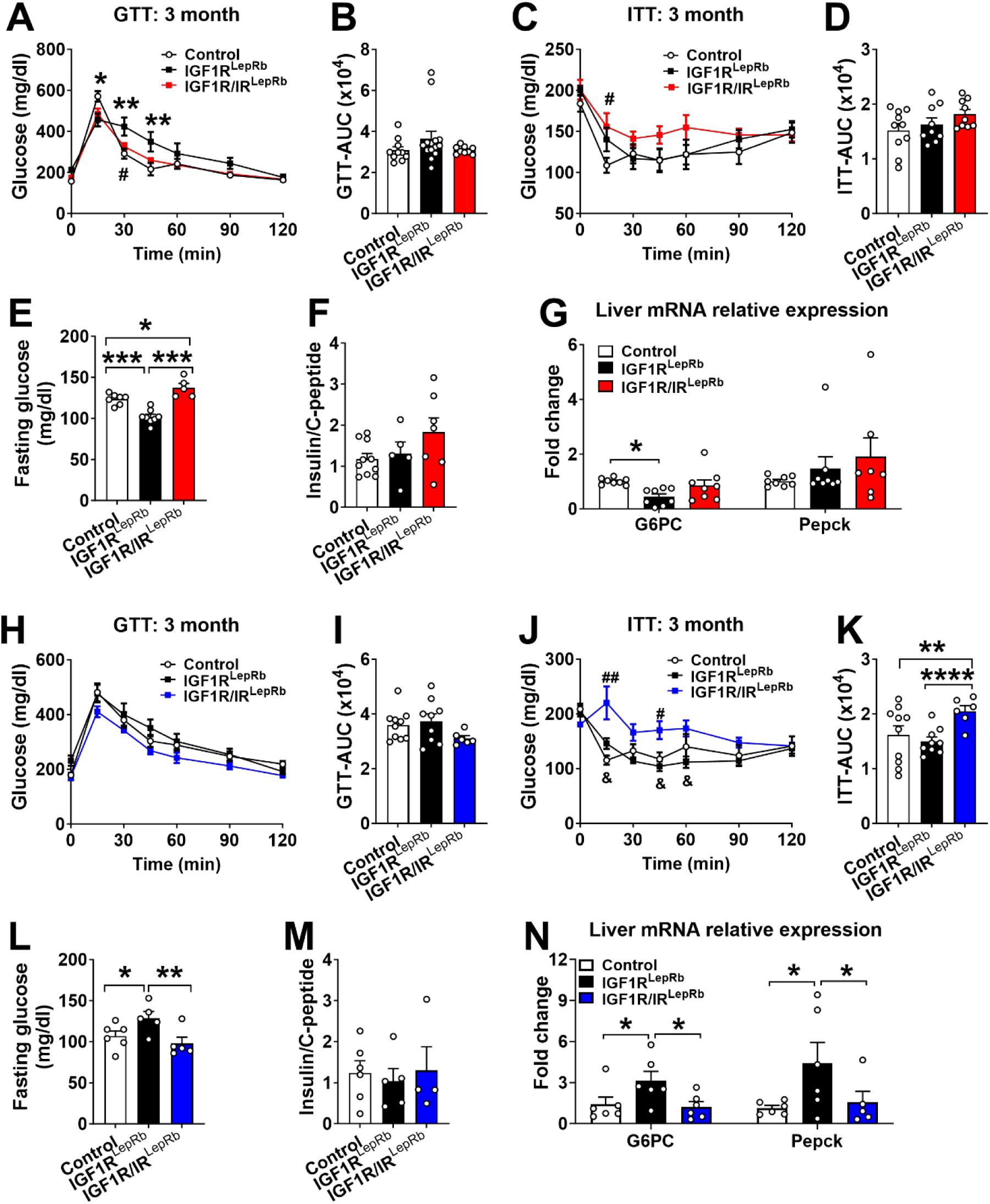
**Changes in glucose homeostasis in IGF1R^LepRb^ and IGF1R/IR^LepRb^ mice**. (A) Glucose tolerance test (GTT), (B) area under the curve of GTT (GTT-AUC), (C) insulin tolerance test (ITT), and 1. (D) AUC of ITT (ITT-AUC) of 3-month-old female control, IGF1R^LepRb^ and IGF1R/IR^LepRb^ mice (n=9-14/group). (E) Fasting glucose and (F) insulin/C-peptide levels in 3-month-old female control, IGF1R^LepRb,^ and IGF1R/IR^LepRb^ mice (n=5-10/group). (G) Relative expression of gluconeogenesis and inflammatory markers in the liver as measured by quantitative PCR in 5-month-old female control, IGF1R^LepRb,^ and IGF1R/IR^LepRb^ mice (n=8/group). (H) GTT, (I) GTT-AUC, (J) ITT, and (K) ITT-AUC of 3-month-old male control, IGF1R^LepRb^ and IGF1R/IR^LepRb^ mice (n=6-10/group). (L) Fasting glucose and (M) insulin/C-peptide levels in 3-month-old male control, IGF1R^LepRb,^ and IGF1R/IR^LepRb^ mice (n=5-6/group). (N) Relative expression of gluconeogenesis markers in the liver as measured by quantitative PCR in 5-month-old male control, IGF1R^LepRb^ and IGF1R/IR^LepRb^ mice (n=6/group). Values throughout the figure are means ±SEM. For the entire figure, \**P* < 0.05, \*\**P* < 0.01, \*\*\**P* < 0.0001, and \*\*\*\**P* < 0.00001, were determined by Bonferroni’s Multiple Comparison Test following one-way ANOVA; or Tukey’s post hoc test following one-way ANOVA.

(Supplemental Fig. 1C) were comparable between female IGF1R^LepRb^ mice and controls. Interestingly, female IGF1R^LepRb^ mice showed decreased fasting glucose (Fig. 8F), suggesting an impaired hepatic gluconeogenic pathway. Then we measured mRNA expression of gene markers in the liver and found gluconeogenic makers were significantly changed including decreased glucose 6-phosphatase (G6PC) and increased phosphoenolpyruvate carboxykinase (Pepck) mRNA expressions (Fig. 8G). Female IGF1R/IR^LepRb^ mice showed insulin insensitivity at 15 min and an increasing trend of ITT-AUC (Fig. 8C-D). Therefore, these results imply that IGF1R in LepRb neurons regulates the hepatic gluconeogenic pathway in female mice to permit normal glucose tolerance and normal fasting glucose.

Male IGF1R^LepRb^ mice did not show glucose intolerance or insulin insensitivity (Fig. 8H-K). In contrast to females, they showed higher fasting glucose levels and elevated mRNA expression of G6PC and Pepck (Fig. 8M-N). Only male IGF1R/IR^LepRb^ mice had insulin insensitivity (Fig. 8J-K), indicating that IGF1R and IR in LepRb neurons jointly regulate insulin sensitivity in male mice. In addition, the altered fasting glucose levels and gluconeogenic genes expression seen in male IGF1R^LepRb^ mice were reversed in male IGF1R/IR^LepRb^ mice (Fig. 8M-N). These results imply that IGF1R and IR play unique and divergent roles in the regulation of glucose homeostasis in both sexes.

## DISCUSSION

This present study showed that the IGF1R in LepRb expressing cells is indispensable for various physiologic processes including reproduction, growth, bone mass accrual, energy balance, and fasting glucose levels with corresponding gluconeogenesis-related gene expressions in the liver (opposite effects seen in females and males) as summarized in Table 2. Previous evidence found that genetic ablation of IR alone in LepRb neurons caused a mild delay in puberty (*11*). Here we further identified that IR and IGF1R jointly regulate many aspects of reproduction, body composition, energy homeostasis, and male insulin sensitivity.

**Table 2.**
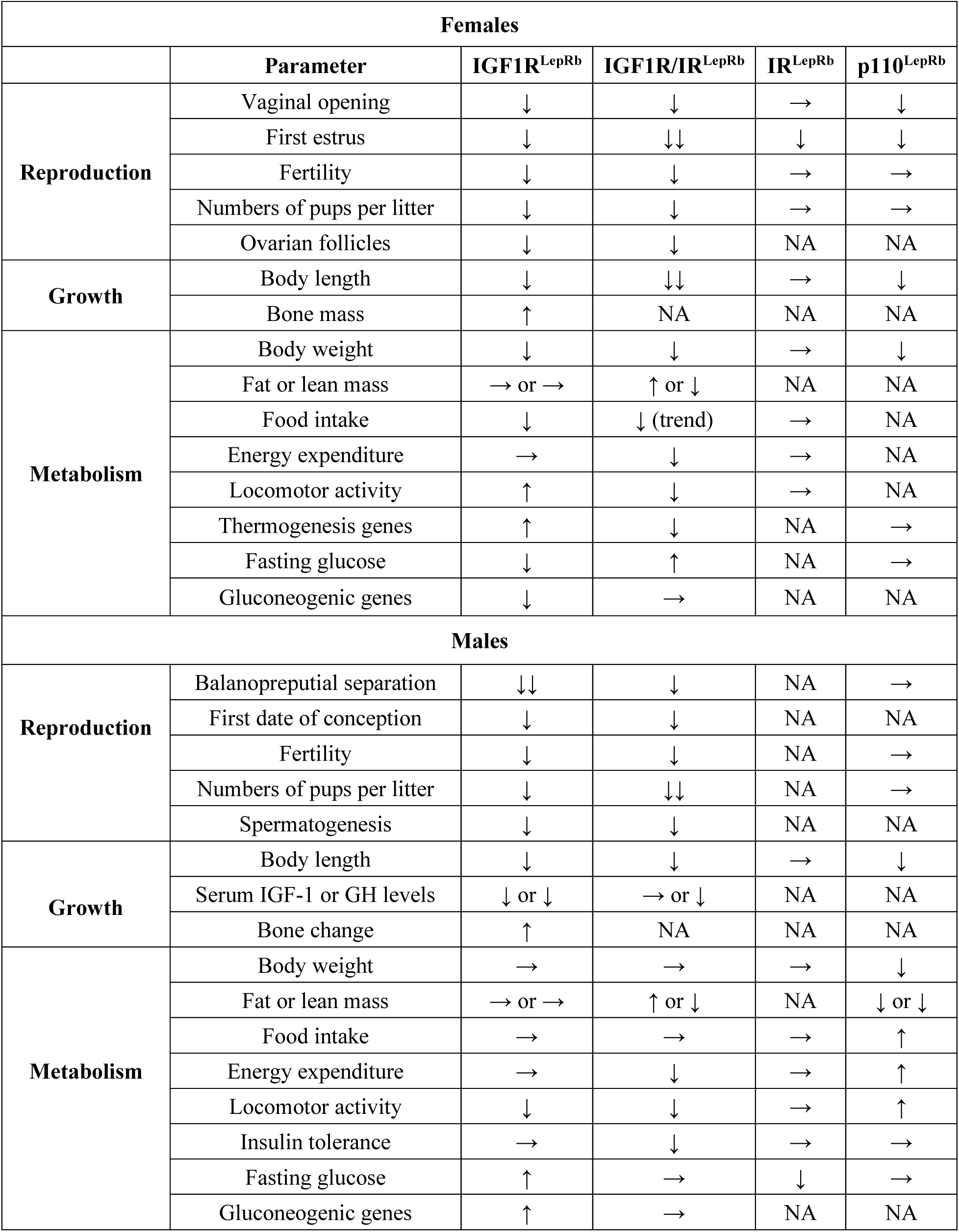
Summary of phenotypic changes in IGF1R^LepRb^ and IGF1R/IR^LepRb^ mice.

| **Females** |  |  |  |  |  |
| --- | --- | --- | --- | --- | --- |
|  | **Parameter** | **IGF1R<sup>LepRb</sup>** | **IGF1R/IR<sup>LepRb</sup>** | **IR<sup>LepRb</sup>** | **p110<sup>LepRb</sup>** |
| **Reproduction** | Vaginal opening | ↓ | ↓ | → | ↓ |
|  | First estrus | ↓ | ↓↓ | ↓ | ↓ |
|  | Fertility | ↓ | ↓ | → | → |
|  | Numbers of pups per litter | ↓ | ↓ | → | → |
|  | Ovarian follicles | ↓ | ↓ | NA | NA |
| **Growth** | Body length | ↓ | ↓↓ | → | ↓ |
|  | Bone mass | ↑ | NA | NA | NA |
| **Metabolism** | Body weight | ↓ | ↓ | → | ↓ |
|  | Fat or lean mass | → or → | ↑ or ↓ | NA | NA |
|  | Food intake | ↓ | ↓ (trend) | → | NA |
|  | Energy expenditure | → | ↓ | → | NA |
|  | Locomotor activity | ↑ | ↓ | → | NA |
|  | Thermogenesis genes | ↑ | ↓ | NA | → |
|  | Fasting glucose | ↓ | ↑ | NA | → |
|  | Gluconeogenic genes | ↓ | → | NA | NA |
| **Males** |  |  |  |  |  |
| **Reproduction** | Balanopreputial separation | ↓↓ | ↓ | NA | → |
|  | First date of conception | ↓ | ↓ | NA | NA |
|  | Fertility | ↓ | ↓ | NA | → |
|  | Numbers of pups per litter | ↓ | ↓↓ | NA | → |
|  | Spermatogenesis | ↓ | ↓ | NA | NA |
| **Growth** | Body length | ↓ | ↓ | → | ↓ |
|  | Serum IGF-1 or GH levels | ↓ or ↓ | → or ↓ | NA | NA |
|  | Bone change | ↑ | NA | NA | NA |
| **Metabolism** | Body weight | → | → | → | ↓ |
|  | Fat or lean mass | → or → | ↑ or ↓ | NA | ↓ or ↓ |
|  | Food intake | → | → | → | ↑ |
|  | Energy expenditure | → | ↓ | → | ↑ |
|  | Locomotor activity | ↓ | ↓ | → | ↑ |
|  | Insulin tolerance | → | ↓ | → | → |
|  | Fasting glucose | ↑ | → | ↓ | → |
|  | Gluconeogenic genes | ↑ | → | NA | NA |

IGF1R in the mammalian brain strongly promotes the development of the somatotropic axis, regulates GH and IGF-1 secretion, and controls the growth of peripheral tissues, glucose metabolism, and energy storage (*5*). Homozygous brain-specific IGF1R knockout mice showed microcephalic, severed growth retardation, infertility, and unexpectedly higher plasma IGF-1 levels, while heterozygous mutants had initially normal but progressively growth retardation from nearly 3 weeks of age onwards (*5*). Interestingly, these heterozygous mutant mice caught up to normal size at around 4 months of age (*5*). These mice had significantly decreased serum levels of IGF-1 at 4 weeks but increased at 8 weeks and continued to be 30%-40% increased throughout adult life (*5*). Hypophysiotropic somatostatin (SST) neurons sense IGF-1 levels (*46*), but ablation of IGF1R and/or growth hormone receptor (GHR) in SST neurons was not sufficient to influence body growth or serum IGF-1 levels (*47*). In another study, we found that loss of IGF1R in kisspeptin (*Kiss1*) neurons (IGF1R^Kiss1^ mice) displayed growth retardation, as evidenced by a shorter body length throughout life (*48*). These findings imply that multiple redundant and compensatory mechanisms may exist to control the somatotropic/growth axis. Our results suggest that IGF1R in LepRb neurons senses IGF-1 signaling to regulate body growth, as indicated by the temporary growth retardation in IGF1R^LepRb^ and IGF1R/IR^LepRb^ mice of both sexes. We further detected decreased IGF-1 and GH levels at 3 months in these male mice, but no changes were seen in females. One previous study showed that disruption of the p110α subunit of PI3K signaling in LepRb positive cells caused growth retardation in females at postnatal day 60 but had normal serum IGF-1 levels (*11*). We speculate that this might be due to the failure to detect the pulsatile secretion of hormones through a single time point measurement. Interestingly, these p110α subunit knockout female mice displayed decreased bone volume and BMD at postnatal day 60 (*11*). However, we observed increased trabecular bone volume and number per unit length, cortical bone area, and bone strength in IGF1R^LepRb^ female and male mice at 5 months of age. The discrepancy between PI3K and IGF-1 signaling in bone morphology may be related to age. Longitudinal studies of changes in bone mass during growth showed that the greatest increases in bone mass occur after puberty between the ages of 12-15 years in girls and 14-17 years in boys (*49*). After reaching a peak in early adulthood between the ages of 25 to 35 years, bone mass declines and raises the risk of osteoporosis and fractures in later life (*50*). The delayed onset of puberty in IGF1R^LepRb^ mice may postpone the bone mass development-peak and subsequent decline, resulting in increased bone volume, strength of midshaft, and resistance to bending compared to our control mice. In support of this notion, in boys with constitutional delay of growth and puberty, bone turnover can be normal, and BMD can increase in a manner similar to healthy children after adjustment for bone age (*51*). However, the association between delayed puberty and bone turnover remains controversial (*52, 53*). Future studies are needed to fully understand this interesting bone phenotype induced by genetic ablation of IGF1R in LepRb neurons.(this sentence may not be needed) An interesting caveat of presented study is a possible combination of the effects of IGF1R signaling in hypothalamus and in bone. Most of skeletal stem cells, which give rise to bone forming osteoblasts and marrow adipocytes, are expressing LepRb, which activity is determined by a pattern of phosphorylation to trigger downstream signaling pathways(*45, 54*). LepRb regulates bone mass and marrow adiposity in a manner specific to sex and skeletal location(*45*). Skeletal stem cells also express IGF1 and IGF1R which locally regulate bone mass and marrow adiposity, and bone regeneration(*55, 56*). Although, there are no clear evidence that the same skeletal stem cell expresses both receptors, it is very possible that the subpopulation of such cells exists and is affected in our model of IGF1R^LepRb^ deficiency. However, inhibition of IGF1 signaling in bone has a negative effect on bone growth and bone mass accrual(*56–58*), which is counterintuitive to presented here phenotype of increased bone mass in IGF1R^LepRb^ mice. This may suggest that IGF1R signaling in LepRb-positive skeletal stem cells is dominated by IGF1R signaling in LepRb-positive neurons. Future clarification of the IGF1R signaling cross talk between brain and bone may benefit better understanding of sexual and developmental nuances of growth and maturation, as we discussed above.

The role of ARH neurons as gatekeepers of female reproduction and energy allocation is well-established (*59–61*). One recent study reported that the brain-derived cellular communication network factor 3 (CCN3) secreted from the ARH *Kiss1* neurons stimulated mouse and human skeletal stem cell activity, increased bone remodeling and fracture repair in young and old mice of both sexes (*62*). This work further showed that lactation stage-specific expression of CCN3 in female ARH *Kiss1* neurons during lactation is a newly identified brain-bone axis evolved to sustain the skeleton in mammalian mothers and offspring (*62*). Interestingly, the Nephroblastoma overexpressed gene (NOV/CCN3) is an adhesive substrate that, in concert with IGF-1, promotes muscle skeletal cell proliferation and survival (*63*). Here we discovered that IGF1R in LepRb neurons is critical in controlling bone homeostasis and reproduction. Considering the overlapped ARH region and the known interaction, we cannot rule out the possibility that IGF1R in LepRb neurons may influence the interaction between IGF-1 and CCN3 to exert multifaceted effects on bone and reproduction.

Reproduction declines with age (*64, 65*), making research into mechanisms that regulate reproductive ageing and preserve reproductive health vital. Pharmacologic blockage of IGF1R signaling can favorably impact lifespan in female mice (*66*). Yet the role of IGF1R signaling in reproductive ageing is unknown. Here we observed that IGF1R in LepRb protects against reproductive decline with age in both female and male mice. Further research is needed to evaluate the impact of IGF1R in LepRb neurons on age-related changes in female and male reproductive systems affecting the ovaries and testes.

We have found that IGF1R in *Kiss1* neurons is crucial for body growth, energy balance, normal timing of pubertal onset, and male reproductive functions (*48*). IGF1R in ARH *Kiss1* neurons may modulate energy balance through communication with pro-opiomelanocortin (POMC) signaling and activation of sympathetic nervous system activity and BAT thermogenesis (*48*). By comparing the IGF1R and/or IR functions in LepRb neurons and *Kiss1* neurons, we found that 1) the multifaceted roles of IGF1R in these two neuronal populations are quite similar, including regulating the normal timing of pubertal onset, fertility, food intake, mild change of body weight, and body length; and 2) IGF1R and IR jointly regulate body composition, energy expenditure, physical activity, and male glucose homeostasis. Interestingly, IGF1R in LepRb neurons is also critical for controlling bone morphology and male serum IGF-1 and GH levels. The wider role of IGF1R in LepRb neurons than *Kiss1* neurons may be attributed to the classical role of leptin signaling in energy balance. One recent study showed that leptin may be a direct effector of linear growth programming by nutrition and that the growth hormone-releasing hormone neuronal subpopulation may display a specific response to leptin in cases of underfeeding (*67*). Leptin acts to upregulate anorexigenic POMC expression (*68, 69*) while downregulating orexigenic agouti-related protein (AgRP) expression and inhibiting AgRP cell activity (*70–72*). POMC neurons innervate the reproductive circuits in the central nervous system and are well-positioned to provide synaptic inputs to GnRH neurons (*73, 74*). Female IGF1RLepRb mice exhibited hypophagia and increased BAT mRNA expressions, consistent with the anorexigenic effects of POMC signaling. Given the importance of the ARH leptin-melanocortin-kisspeptin pathway in the metabolic control of reproduction (*75*), one interesting research direction is to explore the role of IGF1R in POMC neurons.

The lower fasting glucose levels in female IGF1R^LepRb^ mice were associated with decreased hepatic gluconeogenesis as indicated by the lower G6PC expression in the liver. Previous studies have shown that impaired gluconeogenesis-induced hepatic insulin resistance, yet not examined in this current study, was associated with increased body weight gain and diabetes. Thus, the counteracting effects between food intake and glucose levels led to the overall comparable body weight in female IGF1R^LepRb^ mice. Male IGF1R^LepRb^ mice also showed normal body weight but displayed opposite changes in fasting glucose levels due to the increased hepatic gluconeogenesis as indicated by the higher G6PC expression in the liver. Our findings have identified a sex-specific role of IGF-1R in LepRb neurons in regulating glucose levels and associated hepatic gluconeogenesis.

Strikingly, IGF1R/IR^LepRb^ mice of both sexes exhibited dramatically increased fat mass percentage, decreased lean mass percentage, and disrupted male insulin sensitivity compared to IGF1R^LepRb^ mice and controls (Figure 8). The joint role of IGF1R and IR in peripheral tissues, including fat and muscle, were well characterized (*29, 30*). Mice with a combined knockout of IGF1R and IR in fat (FIGIRKO mice) had lower WAT and BAT (*30*), while energy expenditure was higher (*29, 30*). Previous studies have shown that IGF1R and IR play divergent roles in regulating fat mass in the CNS and periphery (*5, 76–78*). IGF1R only modestly contributes to fat mass formation and function, since FIGFRKO mice had a nearly 25% reduction in white fat mass (*76*). In contrast, heterozygous brain IGF1R knockout mice had enlarged fat mass and glucose intolerance (*5*). Similarly, FIRKO mice had a 95% reduction in WAT and are protected against obesity-related glucose intolerance (*78*), while NIRKO mice developed diet-sensitive obesity with increases in body fat, mild insulin resistance, and elevated plasma insulin levels (*77*). In addition, IGF1R^Kiss1^ mice (*48*) or IR^Kiss1^ mice (*37*) did not replicate the glucose intolerance seen in heterozygous brain IGF1R knockout (*5*) or NIRKO mice (*77*). Loss of one signal, either IGF1R or IR activation, may eventually be overcome by other environmental and developmental signals to modulate glucose homeostasis. However simultaneous loss of IGF1R and IR in *Kiss1* neurons disrupted glucose homeostasis (*48*). Here, we have also identified that simultaneous loss of IGF1R and IR in LepRb neurons disrupted insulin sensitivity in male mice. Overall, our findings identified the cooperative role of IGF1R and IR in LepRb neurons in regulating body composition and male insulin insensitivity. Although we characterized a comprehensive set of phenotypes in this current study, one limitation is that we did not further examine the involved mechanisms such as the alterations of leptin and insulin signaling in the brain, liver, and muscle.

In summary, our findings have dissected distinct roles for IGF1R and IR in LepRb neurons in reproduction, body growth, bone health, and metabolism. Loss of IGF1R in LepRb neurons confers resistance to obesity due to increased energy expenditure, showing central IGF1R is obesogenic. These effects diminished in IGF1R/IR^LepRb^ mice due to decreased energy expenditure and physical activity and increased lipid storage in BAT, suggesting IR in LepRb neurons has an overall protective effect against obesity. In addition, our findings provide novel evidence that IGF1R and IR signaling in LepRb neurons coordinate to regulate body composition and insulin sensitivity. These findings extend our understanding of the role of central IGF1R and IR in LepRb neurons in the control of processes including growth, reproduction, body composition, energy balance, and glucose homeostasis.

### AUTHOR CONTRIBUTIONS

Conceptualization: JWH; Data Collection: MW, PC; Formal Analysis: MW, BLC, JWH; Manuscript writing: MW, JWH; Supervision; YX, JWH.

## ACKNOWLEDGEMENT

This work was supported by the National Institutes of Health grant F32HD112123 to MW, HD104418 to JWH, and U.S. Department of Agriculture Grant 51000-064-01S to YX.

---

**Supplemental Figure 1.**
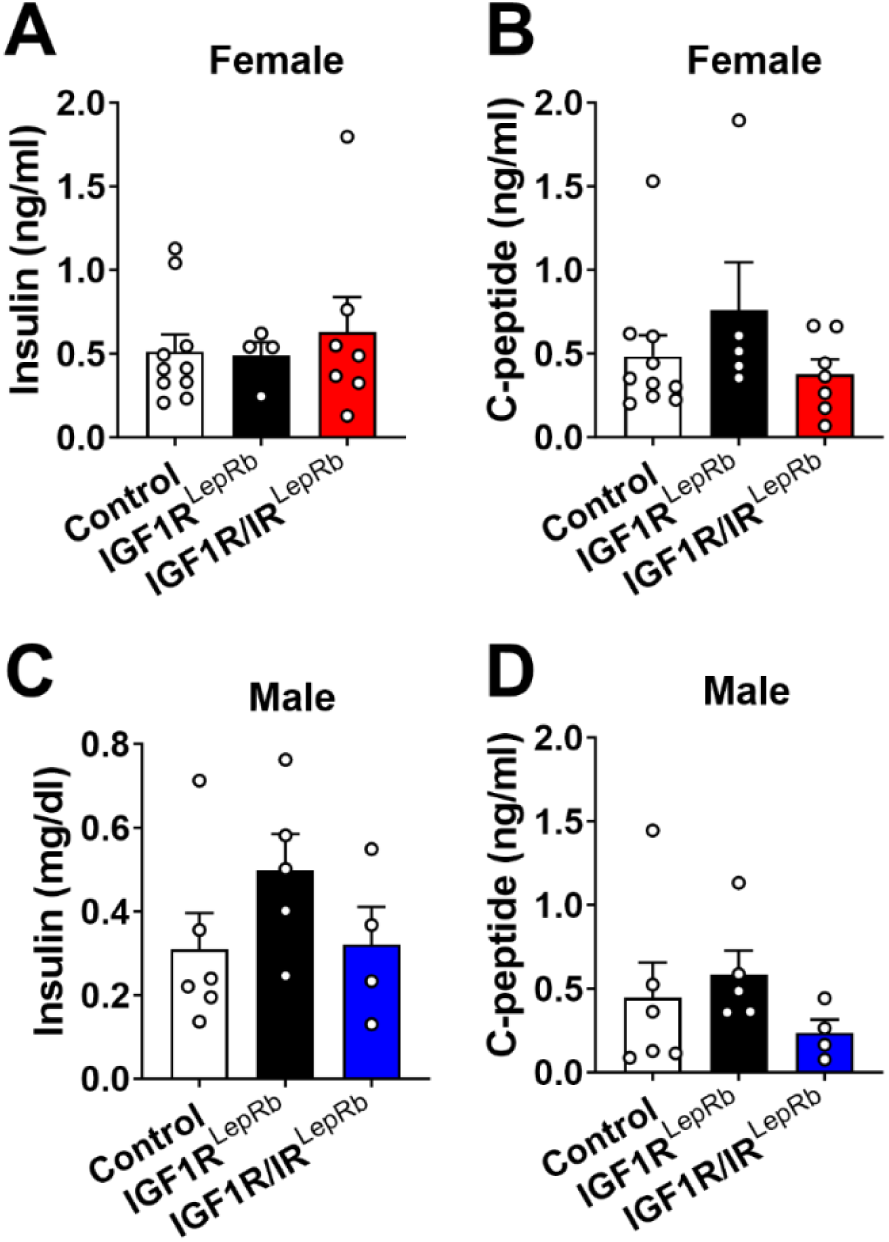
Serum insulin and C-peptide levels in mice. (A and B) Serum insulin and C-peptide in 4-month-old female control, IGF1R^LepRb,^ and IGF1R/IR^LepRb^ mice (n=5-10/group). (C and D) Serum insulin and C-peptide in 4-month-old male control, IGF1R^LepRb,^ and IGF1R/IR^LepRb^ mice (n=4- 6/group). Values throughout figure are means ±SEM. For the entire figure, \**P* < 0.05, determined by t test.

## References

1. D. Garcia-Galiano, B. C. Borges, S. J. Allen, C. F. Elias, PI3K signalling in leptin receptor cells: Role in growth and reproduction. J Neuroendocrinol 31, e12685 (2019).

2. H. Christou, S. Serdy, C. S. Mantzoros, Leptin in relation to growth and developmental processes in the fetus. Semin Reprod Med 20, 123–130 (2002).

3. D. Le Roith, C. Bondy, S. Yakar, J. L. Liu, A. Butler, The somatomedin hypothesis: 2001. Endocr Rev 22, 53–74 (2001).

4. S. A. Divall, T. R. Williams, S. E. Carver, L. Koch, J. C. Bruning, C. R. Kahn, F. Wondisford, S. Radovick, A. Wolfe, Divergent roles of growth factors in the GnRH regulation of puberty in mice. The Journal of clinical investigation 120, 2900–2909 (2010).

5. L. Kappeler, C. De Magalhaes Filho, J. Dupont, P. Leneuve, P. Cervera, L. Perin, C. Loudes, A. Blaise, R. Klein, J. Epelbaum, Y. Le Bouc, M. Holzenberger, Brain IGF-1 receptors control mammalian growth and lifespan through a neuroendocrine mechanism. PLoS Biol 6, e254 (2008).

6. A. S. Banks, S. M. Davis, S. H. Bates, M. G. Myers, Jr., Activation of downstream signals by the long form of the leptin receptor. The Journal of biological chemistry 275, 14563–14572 (2000).

7. W. A. Banks, Leptin transport across the blood-brain barrier: implications for the cause and treatment of obesity. Current pharmaceutical design 7, 125–133 (2001).

8. S. C. Chua, Jr., S. M. Liu, Q. Li, A. Sun, W. F. DeNino, S. B. Heymsfield, X. E. Guo, Transgenic complementation of leptin receptor deficiency. II. Increased leptin receptor transgene dose effects on obesity/diabetes and fertility/lactation in lepr-db/db mice. American journal of physiology. Endocrinology and metabolism 286, E384–392 (2004).

9. C. de Luca, T. J. Kowalski, Y. Zhang, J. K. Elmquist, C. Lee, M. W. Kilimann, T. Ludwig, S. M. Liu, S. C. Chua, Jr., Complete rescue of obesity, diabetes, and infertility in db/db mice by neuron-specific LEPR-B transgenes. The Journal of clinical investigation 115, 3484–3493 (2005).

10. S. A. Robertson, G. M. Leinninger, M. G. Myers, Jr., Molecular and neural mediators of leptin action. Physiology & behavior 94, 637–642 (2008).

11. D. Garcia-Galiano, B. C. Borges, J. Donato, Jr., S. J. Allen, N. Bellefontaine, M. Wang, J. J. Zhao, K. M. Kozloff, J. W. Hill, C. F. Elias, PI3Kalpha inactivation in leptin receptor cells increases leptin sensitivity but disrupts growth and reproduction. JCI insight 2, (2017).

12. M. Sadagurski, R. L. Leshan, C. Patterson, A. Rozzo, A. Kuznetsova, J. Skorupski, J. C. Jones, R. A. Depinho, M. G. Myers, Jr., M. F. White, IRS2 signaling in LepR-b neurons suppresses FoxO1 to control energy balance independently of leptin action. Cell metabolism 15, 703–712 (2012).

13. L. Plum, E. Rother, H. Munzberg, F. T. Wunderlich, D. A. Morgan, B. Hampel, M. Shanabrough, R. Janoschek, A. C. Konner, J. Alber, A. Suzuki, W. Krone, T. L. Horvath, K. Rahmouni, J. C. Bruning, Enhanced leptin-stimulated Pi3k activation in the CNS promotes white adipose tissue transdifferentiation. Cell metabolism 6, 431–445 (2007).

14. M. Bjornholm, H. Munzberg, R. L. Leshan, E. C. Villanueva, S. H. Bates, G. W. Louis, J. C. Jones, R. Ishida-Takahashi, C. Bjorbaek, M. G. Myers, Jr., Mice lacking inhibitory leptin receptor signals are lean with normal endocrine function. The Journal of clinical investigation 117, 1354–1360 (2007).

15. C. M. Patterson, E. C. Villanueva, M. Greenwald-Yarnell, M. Rajala, I. E. Gonzalez, N. Saini, J. Jones, M. G. Myers, Jr., Leptin action via LepR-b Tyr1077 contributes to the control of energy balance and female reproduction. Molecular metabolism 1, 61–69 (2012).

16. B. C. Borges, D. Garcia-Galiano, R. Rorato, L. L. K. Elias, C. F. Elias, PI3K p110beta subunit in leptin receptor expressing cells is required for the acute hypophagia induced by endotoxemia. Molecular metabolism 5, 379–391 (2016).

17. K. J. Motyl, C. J. Rosen, Understanding leptin-dependent regulation of skeletal homeostasis. Biochimie 94, 2089–2096 (2012).

18. H. Liu, Y. He, J. Bai, C. Zhang, F. Zhang, Y. Yang, H. Luo, M. Yu, H. Liu, L. Tu, N. Zhang, N. Yin, J. Han, Z. Yan, N. A. Scarcelli, K. M. Conde, M. Wang, J. C. Bean, C. H. S. Potts, C. Wang, F. Hu, F. Liu, Y. Xu, Hypothalamic Grb10 enhances leptin signalling and promotes weight loss. Nat Metab 5, 147–164 (2023).

19. J. K. Elmquist, C. F. Elias, C. B. Saper, From lesions to leptin: hypothalamic control of food intake and body weight. Neuron 22, 221–232 (1999).

20. M. W. Schwartz, S. C. Woods, D. Porte, Jr., R. J. Seeley, D. G. Baskin, Central nervous system control of food intake. Nature 404, 661–671 (2000).

21. E. van de Wall, R. Leshan, A. W. Xu, N. Balthasar, R. Coppari, S. M. Liu, Y. H. Jo, R. G. MacKenzie, D. B. Allison, N. J. Dun, J. Elmquist, B. B. Lowell, G. S. Barsh, C. de Luca, M. G. Myers, Jr., G. J. Schwartz, S. C. Chua, Jr., Collective and individual functions of leptin receptor modulated neurons controlling metabolism and ingestion. Endocrinology 149, 1773–1785 (2008).

22. N. Balthasar, R. Coppari, J. McMinn, S. M. Liu, C. E. Lee, V. Tang, C. D. Kenny, R. A. McGovern, S. C. Chua, Jr., J. K. Elmquist, B. B. Lowell, Leptin receptor signaling in POMC neurons is required for normal body weight homeostasis. Neuron 42, 983–991 (2004).

23. A. C. Rupp, A. J. Tomlinson, A. H. Affinati, W. T. Yacawych, A. M. Duensing, C. True, S. R. Lindsley, M. A. Kirigiti, A. MacKenzie, J. Polex-Wolf, C. Li, L. B. Knudsen, R. J. Seeley, D. P. Olson, P. Kievit, M. G. Myers, Jr., Suppression of food intake by Glp1r/Lepr-coexpressing neurons prevents obesity in mouse models. The Journal of clinical investigation 133, (2023).

24. Y. Zhang, I. A. Kerman, A. Laque, P. Nguyen, M. Faouzi, G. W. Louis, J. C. Jones, C. Rhodes, H. Munzberg, Leptin-receptor-expressing neurons in the dorsomedial hypothalamus and median preoptic area regulate sympathetic brown adipose tissue circuits. J Neurosci 31, 1873–1884 (2011).

25. G. Cady, T. Landeryou, M. Garratt, J. J. Kopchick, N. Qi, D. Garcia-Galiano, C. F. Elias, M. G. Myers, Jr., R. A. Miller, D. A. Sandoval, M. Sadagurski, Hypothalamic growth hormone receptor (GHR) controls hepatic glucose production in nutrient-sensing leptin receptor (LepRb) expressing neurons. Molecular metabolism 6, 393–405 (2017).

26. A. M. Valverde, D. J. Burks, I. Fabregat, T. L. Fisher, J. Carretero, M. F. White, M. Benito, Molecular mechanisms of insulin resistance in IRS-2-deficient hepatocytes. Diabetes 52, 2239–2248 (2003).

27. M. Matsumoto, D. Accili, All roads lead to FoxO. Cell metabolism 1, 215–216 (2005).

28. A. Taguchi, L. M. Wartschow, M. F. White, Brain IRS2 signaling coordinates life span and nutrient homeostasis. Science 317, 369–372 (2007).

29. B. T. O’Neill, H. P. Lauritzen, M. F. Hirshman, G. Smyth, L. J. Goodyear, C. R. Kahn, Differential Role of Insulin/IGF-1 Receptor Signaling in Muscle Growth and Glucose Homeostasis. Cell reports 11, 1220–1235 (2015).

30. J. Boucher, M. A. Mori, K. Y. Lee, G. Smyth, C. W. Liew, Y. Macotela, M. Rourk, M. Bluher, S. J. Russell, C. R. Kahn, Impaired thermogenesis and adipose tissue development in mice with fat-specific disruption of insulin and IGF-1 signalling. Nat Commun 3, 902 (2012).

31. J. DeFalco, M. Tomishima, H. Liu, C. Zhao, X. Cai, J. D. Marth, L. Enquist, J. M. Friedman, Virus-assisted mapping of neural inputs to a feeding center in the hypothalamus. Science 291, 2608–2613 (2001).

32. N. Kloting, L. Koch, T. Wunderlich, M. Kern, K. Ruschke, W. Krone, J. C. Bruning, M. Bluher, Autocrine IGF-1 action in adipocytes controls systemic IGF-1 concentrations and growth. Diabetes 57, 2074–2082 (2008).

33. H. Stachelscheid, H. Ibrahim, L. Koch, A. Schmitz, M. Tscharntke, F. T. Wunderlich, J. Scott, C. Michels, C. Wickenhauser, I. Haase, J. C. Bruning, C. M. Niessen, Epidermal insulin/IGF-1 signalling control interfollicular morphogenesis and proliferative potential through Rac activation. The EMBO journal 27, 2091–2101 (2008).

34. J. C. Bruning, M. D. Michael, J. N. Winnay, T. Hayashi, D. Horsch, D. Accili, L. J. Goodyear, C. R. Kahn, A muscle-specific insulin receptor knockout exhibits features of the metabolic syndrome of NIDDM without altering glucose tolerance. Molecular cell 2, 559–569 (1998).

35. L. Madisen, T. A. Zwingman, S. M. Sunkin, S. W. Oh, H. A. Zariwala, H. Gu, L. L. Ng, R. D. Palmiter, M. J. Hawrylycz, A. R. Jones, E. S. Lein, H. Zeng, A robust and high-throughput Cre reporting and characterization system for the whole mouse brain. Nature neuroscience 13, 133–140 (2010).

36. M. A. Torsoni, B. C. Borges, J. L. Cote, S. J. Allen, E. Mahany, D. Garcia-Galiano, C. F. Elias, AMPKalpha2 in Kiss1 Neurons Is Required for Reproductive Adaptations to Acute Metabolic Challenges in Adult Female Mice. Endocrinology 157, 4803–4816 (2016).

37. X. Qiu, A. R. Dowling, J. S. Marino, L. D. Faulkner, B. Bryant, J. C. Bruning, C. F. Elias, J. W. Hill, Delayed puberty but normal fertility in mice with selective deletion of insulin receptors from Kiss1 cells. Endocrinology 154, 1337–1348 (2013).

38. J. W. Hill, Y. Xu, F. Preitner, M. Fukuda, Y. R. Cho, J. Luo, N. Balthasar, R. Coppari, L. C. Cantley, B. B. Kahn, J. J. Zhao, J. K. Elmquist, Phosphatidyl inositol 3-kinase signaling in hypothalamic proopiomelanocortin neurons contributes to the regulation of glucose homeostasis. Endocrinology 150, 4874–4882 (2009).

39. A. Mesaros, S. B. Koralov, E. Rother, F. T. Wunderlich, M. B. Ernst, G. S. Barsh, K. Rajewsky, J. C. Bruning, Activation of Stat3 signaling in AgRP neurons promotes locomotor activity. Cell metabolism 7, 236–248 (2008).

40. L. A. Stechschulte, P. J. Czernik, Z. C. Rotter, F. N. Tausif, C. A. Corzo, D. P. Marciano, A. Asteian, J. Zheng, J. B. Bruning, T. M. Kamenecka, C. J. Rosen, P. R. Griffin, B. Lecka-Czernik, PPARG Post-translational Modifications Regulate Bone Formation and Bone Resorption. EBioMedicine 10, 174–184 (2016).

41. M. L. Bouxsein, S. K. Boyd, B. A. Christiansen, R. E. Guldberg, K. J. Jepsen, R. Muller, Guidelines for assessment of bone microstructure in rodents using micro-computed tomography. Journal of bone and mineral research : the official journal of the American Society for Bone and Mineral Research 25, 1468–1486 (2010).

42. Z. Shi, H. Enayatullah, Z. Lv, H. Dai, Q. Wei, L. Shen, B. Karwand, F. Shi, Freeze-Dried Royal Jelly Proteins Enhanced the Testicular Development and Spermatogenesis in Pubescent Male Mice. Animals (Basel) 9, (2019).

43. B. Lecka-Czernik, L. A. Stechschulte, P. J. Czernik, S. B. Sherman, S. Huang, A. Krings, Marrow Adipose Tissue: Skeletal Location, Sexual Dimorphism, and Response to Sex Steroid Deficiency. Frontiers in endocrinology 8, 188 (2017).

44. F. J. C. Dolcetti, E. Falomir-Lockhart, F. Acuna, M. L. Herrera, S. Cervellini, C. G. Barbeito, D. Grassi, M. A. Arevalo, M. J. Bellini, IGF1 gene therapy in middle-aged female rats delays reproductive senescence through its effects on hypothalamic GnRH and kisspeptin neurons. Aging (Albany NY) 14, 8615–8632 (2022).

45. I. C. McCabe, A. Fedorko, M. G. Myers, Jr., G. Leinninger, E. Scheller, L. R. McCabe, Novel leptin receptor signaling mutants identify location and sex-dependent modulation of bone density, adiposity, and growth. J Cell Biochem 120, 4398–4408 (2019).

46. M. Sato, L. A. Frohman, Differential effects of central and peripheral administration of growth hormone (GH) and insulin-like growth factor on hypothalamic GH-releasing hormone and somatostatin gene expression in GH-deficient dwarf rats. Endocrinology 133, 793–799 (1993).

47. F. M. Chaves, F. Wasinski, M. R. Tavares, N. S. Mansano, R. Frazao, D. O. Gusmao, P. G. F. Quaresma, J. A. B. Pedroso, C. F. Elias, E. O. List, J. J. Kopchick, R. E. Szawka, J. Donato, Effects of the Isolated and Combined Ablation of Growth Hormone and IGF-1 Receptors in Somatostatin Neurons. Endocrinology 163, (2022).

48. M. Wang, S. M. Pugh, J. Daboul, D. Miller, Y. Xu, J. W. Hill, IGF-1 Acts through Kiss1-expressing Cells to Influence Metabolism and Reproduction. bioRxiv, (2024).

49. G. Theintz, B. Buchs, R. Rizzoli, D. Slosman, H. Clavien, P. C. Sizonenko, J. P. Bonjour, Longitudinal monitoring of bone mass accumulation in healthy adolescents: evidence for a marked reduction after 16 years of age at the levels of lumbar spine and femoral neck in female subjects. The Journal of clinical endocrinology and metabolism 75, 1060–1065 (1992).

50. S. H. Ralston, The genetics of osteoporosis. QJM : monthly journal of the Association of Physicians 90, 247–251 (1997).

51. B. Krupa, T. Miazgowski, Bone mineral density and markers of bone turnover in boys with constitutional delay of growth and puberty. The Journal of clinical endocrinology and metabolism 90, 2828–2830 (2005).

52. D. L. Cousminer, J. A. Mitchell, A. Chesi, S. M. Roy, H. J. Kalkwarf, J. M. Lappe, V. Gilsanz, S. E. Oberfield, J. A. Shepherd, A. Kelly, S. E. McCormack, B. F. Voight, B. S. Zemel, S. F. Grant, Genetically Determined Later Puberty Impacts Lowered Bone Mineral Density in Childhood and Adulthood. Journal of bone and mineral research : the official journal of the American Society for Bone and Mineral Research 33, 430–436 (2018).

53. A. Elhakeem, M. Frysz, K. Tilling, J. H. Tobias, D. A. Lawlor, Association Between Age at Puberty and Bone Accrual From 10 to 25 Years of Age. JAMA Netw Open 2, e198918 (2019).

54. B. O. Zhou, R. Yue, M. M. Murphy, J. G. Peyer, S. J. Morrison, Leptin-receptor-expressing mesenchymal stromal cells represent the main source of bone formed by adult bone marrow. Cell Stem Cell 15, 154–168 (2014).

55. B. Lecka-Czernik, C. Ackert-Bicknell, M. L. Adamo, V. Marmolejos, G. A. Churchill, K. R. Shockley, I. R. Reid, A. Grey, C. J. Rosen, Activation of peroxisome proliferator-activated receptor gamma (PPARgamma) by rosiglitazone suppresses components of the insulin-like growth factor regulatory system in vitro and in vivo. Endocrinology 148, 903–911 (2007).

56. J. Wang, Q. Zhu, D. Cao, Q. Peng, X. Zhang, C. Li, C. Zhang, B. O. Zhou, R. Yue, Bone marrow-derived IGF-1 orchestrates maintenance and regeneration of the adult skeleton. Proc Natl Acad Sci U S A 120, e2203779120 (2023).

57. H. L. Racine, M. A. Serrat, The Actions of IGF-1 in the Growth Plate and Its Role in Postnatal Bone Elongation. Curr Osteoporos Rep 18, 210–227 (2020).

58. M. Zhang, S. Xuan, M. L. Bouxsein, D. von Stechow, N. Akeno, M. C. Faugere, H. Malluche, G. Zhao, C. J. Rosen, A. Efstratiadis, T. L. Clemens, Osteoblast-specific knockout of the insulin-like growth factor (IGF) receptor gene reveals an essential role of IGF signaling in bone matrix mineralization. The Journal of biological chemistry 277, 44005–44012 (2002).

59. V. M. Navarro, Metabolic regulation of kisspeptin - the link between energy balance and reproduction. Nat Rev Endocrinol 16, 407–420 (2020).

60. S. L. Padilla, J. G. Perez, M. Ben-Hamo, C. W. Johnson, R. E. A. Sanchez, I. L. Bussi, R. D. Palmiter, H. O. de la Iglesia, Kisspeptin Neurons in the Arcuate Nucleus of the Hypothalamus Orchestrate Circadian Rhythms and Metabolism. Curr Biol 29, 592–604 e594 (2019).

61. L. Wang, C. Vanacker, L. L. Burger, T. Barnes, Y. M. Shah, M. G. Myers, S. M. Moenter, Genetic dissection of the different roles of hypothalamic kisspeptin neurons in regulating female reproduction. Elife 8, (2019).

62. M. E. Babey, W. C. Krause, K. Chen, C. B. Herber, Z. Torok, J. Nikkanen, R. Rodriguez, X. Zhang, F. Castro-Navarro, Y. Wang, E. E. Wheeler, S. Villeda, J. K. Leach, N. E. Lane, E. L. Scheller, C. K. F. Chan, T. H. Ambrosi, H. A. Ingraham, A maternal brain hormone that builds bone. Nature, (2024).

63. J. Lafont, H. Thibout, C. Dubois, M. Laurent, C. Martinerie, NOV/CCN3 induces adhesion of muscle skeletal cells and cooperates with FGF2 and IGF-1 to promote proliferation and survival. Cell Commun Adhes 12, 41–57 (2005).

64. J. A. Martin, B. E. Hamilton, M. J. K. Osterman, A. K. Driscoll, Births: Final Data for 2019. Natl Vital Stat Rep 70, 1–51 (2021).

65. A. Owen, K. Carlson, P. B. Sparzak, "Age-Related Fertility Decline" in StatPearls (Treasure Island (FL), 2024).

66. K. Mao, G. F. Quipildor, T. Tabrizian, A. Novaj, F. Guan, R. O. Walters, F. Delahaye, G. B. Hubbard, Y. Ikeno, K. Ejima, P. Li, D. B. Allison, H. Salimi-Moosavi, P. J. Beltran, P. Cohen, N. Barzilai, D. M. Huffman, Late-life targeting of the IGF-1 receptor improves healthspan and lifespan in female mice. Nat Commun 9, 2394 (2018).

67. L. Decourtye-Espiard, M. Clemessy, P. Leneuve, E. Mire, T. Ledent, Y. Le Bouc, L. Kappeler, Stimulation of GHRH Neuron Axon Growth by Leptin and Impact of Nutrition during Suckling in Mice. Nutrients 15, (2023).

68. M. W. Schwartz, R. J. Seeley, S. C. Woods, D. S. Weigle, L. A. Campfield, P. Burn, D. G. Baskin, Leptin increases hypothalamic pro-opiomelanocortin mRNA expression in the rostral arcuate nucleus. Diabetes 46, 2119–2123 (1997).

69. J. E. Thornton, C. C. Cheung, D. K. Clifton, R. A. Steiner, Regulation of hypothalamic proopiomelanocortin mRNA by leptin in ob/ob mice. Endocrinology 138, 5063–5066 (1997).

70. J. P. Wilding, S. G. Gilbey, C. J. Bailey, R. A. Batt, G. Williams, M. A. Ghatei, S. R. Bloom, Increased neuropeptide-Y messenger ribonucleic acid (mRNA) and decreased neurotensin mRNA in the hypothalamus of the obese (ob/ob) mouse. Endocrinology 132, 1939–1944 (1993).

71. M. W. Schwartz, R. J. Seeley, L. A. Campfield, P. Burn, D. G. Baskin, Identification of targets of leptin action in rat hypothalamus. The Journal of clinical investigation 98, 1101–1106 (1996).

72. C. F. Elias, C. Aschkenasi, C. Lee, J. Kelly, R. S. Ahima, C. Bjorbaek, J. S. Flier, C. B. Saper, J. K. Elmquist, Leptin differentially regulates NPY and POMC neurons projecting to the lateral hypothalamic area. Neuron 23, 775–786 (1999).

73. C. Leranth, N. J. MacLusky, M. Shanabrough, F. Naftolin, Immunohistochemical evidence for synaptic connections between pro-opiomelanocortin-immunoreactive axons and LH-RH neurons in the preoptic area of the rat. Brain research 449, 167–176 (1988).

74. F. Pimpinelli, M. Parenti, F. Guzzi, F. Piva, T. Hokfelt, R. Maggi, Presence of delta opioid receptors on a subset of hypothalamic gonadotropin releasing hormone (GnRH) neurons. Brain research 1070, 15–23 (2006).

75. M. Manfredi-Lozano, J. Roa, F. Ruiz-Pino, R. Piet, D. Garcia-Galiano, R. Pineda, A. Zamora, S. Leon, M. A. Sanchez-Garrido, A. Romero-Ruiz, C. Dieguez, M. J. Vazquez, A. E. Herbison, L. Pinilla, M. Tena-Sempere, Defining a novel leptin-melanocortin-kisspeptin pathway involved in the metabolic control of puberty. Molecular metabolism 5, 844–857 (2016).

76. J. Boucher, S. Softic, A. El Ouaamari, M. T. Krumpoch, A. Kleinridders, R. N. Kulkarni, B. T. O’Neill, C. R. Kahn, Differential Roles of Insulin and IGF-1 Receptors in Adipose Tissue Development and Function. Diabetes 65, 2201–2213 (2016).

77. J. C. Bruning, D. Gautam, D. J. Burks, J. Gillette, M. Schubert, P. C. Orban, R. Klein, W. Krone, D. Muller-Wieland, C. R. Kahn, Role of brain insulin receptor in control of body weight and reproduction. Science 289, 2122–2125 (2000).

78. M. Bluher, M. D. Michael, O. D. Peroni, K. Ueki, N. Carter, B. B. Kahn, C. R. Kahn, Adipose tissue selective insulin receptor knockout protects against obesity and obesity-related glucose intolerance. Dev Cell 3, 25–38 (2002).

